# Cationic amino acid identity and net charge influence condensate properties in *E. coli*

**DOI:** 10.1101/2025.03.29.646054

**Authors:** Aaron K. Kidane, Jacob R. Rosenfeld, Jake D. Johnston, Charlie Dubbeldam, Mohammadreza Paraan, Allie C. Obermeyer

## Abstract

Understanding the formation of biomolecular condensates (BMC) in biological systems has proven to be a paradigm shift in our understanding of the subcellular organization of biomacromolecules. From RNA metabolism, stress response mechanisms, and amyloidogenic pathologies, condensates have been implicated to play a role in a myriad of cellular phenomena. Despite their near ubiquity, we still do no wholly understand how the primary sequence of biomolecules influences their biophysical and rheological properties. Here, we aim to understand the impact of primary cationic amino acid composition on the properties of condensates. Using engineered recombinant proteins, we show that the formation and phase boundaries of coacervates formed between proteins and RNA is dependent on the cationic amino acid identity, as well as the net charge of the protein involved in condensation. Despite the equivalent charge between arginine and lysine at physiological pH, arginine has been shown to promote increased encapsulation efficiency and salt stability, as well as reduced protein mobility within condensates. We show that arginine-tagged globular proteins also have a higher salt resistance *in vitro* when compared to similar lysine-tagged globular proteins. This translates to a cellular context in which arginine tagged proteins promote increased condensate formation in model *E. coli* cells. We were also able to observe a reduction in the total fluorescent recovery and protein mobility within arginine-based condensates via FRAP. Together, these results suggest that in addition to electrostatic interactions and disorder as the main driving forces of phase separation in biological contexts, the primary sequence and side chain composition of proteins plays a significant role in dictating dynamics of coacervates.

## Introduction

The formation of biomolecular condensates (BMC) in biological systems has garnered significant interest and scrutiny in recent years. Over the past two decades, understanding of the vital roles condensates play in a myriad of cellular processes has increased rapidly. First proposed in the early 1900’s^1^, the demixing of biomolecules gained new focus nearly 100 years later with the discovery of germline P granules in *C. elegans*^2^. Dynamic condensate formation via phase separation has been found to play a role in many other cellular processes, not limited to cell-to-cell signal transduction,^3^ heat and osmotic stress-responses^4–6^, gene regulation and RNA metabolism,^7–10^ and heterochromatin packing in the nucleus.^11–14^ In addition, aberrant BMC formation has been found to play a crucial role in numerous neurodegenerative diseases,^15–19^ viral life cycles,^20^ and cancer pathologies.^21–23^ From an engineering perspective, phase separated condensates have shown incredible potential to be utilized for the development of novel protein drug delivery systems, which exhibit high delivery yields in different cell cultures^24,25^. Engineered pH-sensitive coacervates have been developed that dissolve as glucose is metabolized into gluconic acid by glucose oxidase within the dense phase, leading to the controlled release of insulin.^26^ Similarly, negative feedback loops using coacervates and pH-responsive condensation-dissolution cycles with the enzyme catalase have also been recently developed.^27^

Despite the pervasiveness of condensates in cellular physiology, disease pathologies, and engineering applications, only recently have the primary sequence determinants of phase separation begun to be elucidated, as well as the impact of these sequences on the resulting condensates. Currently, our detailed understanding shows that increased valency and biopolymer length, decreased structure and complexity, and the prevalence of charged or aromatic patches contribute to increased phase separation and encapsulation efficiency.^28–32^. Furthermore, the post translational modification (PTM) of charged amino acids has been shown to modulate the net charge and degree of disorder for phase separating proteins^33^, leading to macroscopic changes in condensate behavior. Changes in the cationic amino acid identity of synthetic polypetides in coacervates with RNA has also been shown to have drastic changes on the rheological properties of the resultant condensates^31,34,35^. Polyarginine-based droplets were shown to not only be up to 100 times more viscous than polylysine-based droplets of the same length and charge, but also to outcompete polylysine for in vitro complexation with UTP to form condensates^34^. This difference in behavior is due, in large part, to the resonance states of arginine’s side chain that allows for higher order electrostatic interactions relative to lysine. The delocalization of arginine’s +1 charge on the guanidinium group results in the ability to engage in cation-anion, cation-*π*, and *π*-*π* interactions, whereas the amine group of lysine can only engage in cation-anion and cation-*π* interactions.^28,29^. Complementary studies of intrinsically disordered proteins (IDPs), key to the formation of endogenous BMCs, have also shown that substitution of arginine with lysine results in a dramatic increase in the C_sat_^28^, indicating that arginine residues are preferred for BMC formation due to the myriad of non-covalent interactions.

Here, we aim to understand the impact of variable cationic amino acid identity on the behavior of condensates within the cellular context. Using a panel of four engineered proteins and *E. coli* cells as a model cell, we study the impact of varying cationic side chain composition on the rheological properties of engineered BMCs. The effect of sequence identity on BMC formation has often been studied in relation to its complexation with synthetic polymers, other proteins, or synthetic nucleic acid repeats (PolyU, PolyA, etc.)^31,32,34,36–40^. The characteristics of these cationic side chains have sparsely been investigated in complex with endogenous RNA, so we aim to discern the role of RNA in modulating any observed differences between arginine and lysine. We find that arginine-tagged globular proteins have increased phase boundaries when complexed with RNA, in addition to an increased resistance to salt when compared to isoelectric lysine-based globular proteins. Furthermore, arginine-tagged proteins exogenously expressed in *E. coli* exhibit a much higher partitioning of protein in the condensed phase, and have decreased protein mobility within the dense phase.

## Results and Discussion

### Design of protein variants to probe molecular grammar of engineered condensates

In order to probe the impact of cationic amino acid identity on the phase boundaries in a cellular context, we designed positively charged proteins that were likely to undergo complex coacervation with endogenous RNA. Prior work has shown that *in vitro*, as well as within *E. coli*, proteins with a net charge greater than or equal to +6 have the ability to phase separate with RNA. With this in mind, we expressed and purified a panel of six globular proteins with important variations in their net charge and cationic amino acid composition. We have worked with charge variants of the protein sfGFP, due to its robust fluorescence and high tolerance for engineered mutations^30,41^. GFP(0) and GFP(+6) were created using point mutations on the surface of sfGFP^42^ that shift the net charge at physiological pH from -7 to the ones denoted in the names (SI Table 1). Using these two globular proteins as our starting point, we genetically encoded an Arg-Arg-Arg-Arg-Arg-Arg (6Arg) or Lys-Lys-Lys-Lys-Lys-Lys (6Lys) tag on the C terminus of both globular proteins^41^ (Fig. 1a), creating GFP(+6)-6Arg, GFP(+6)-6Lys, GFP(0)-6Arg, and GFP(0)-6Lys. This panel of 6 proteins with net charges ranging from 0 to +12 lies at or near the charge threshold for complex coacervation with RNA in *E. coli* ^42^, therefore experiments conducted at this charge range allows us to more clearly observe small changes in phase behavior and rheological properties between arginine and lysine containing sequences. All engineered proteins were purified from NiCo21 cells (NEB) using Ni-NTA chromatography, after which the identity and purity were confirmed by MALDI-TOF and SDS-PAGE analysis, respectively (SI Figure 1).

**Figure 1:**
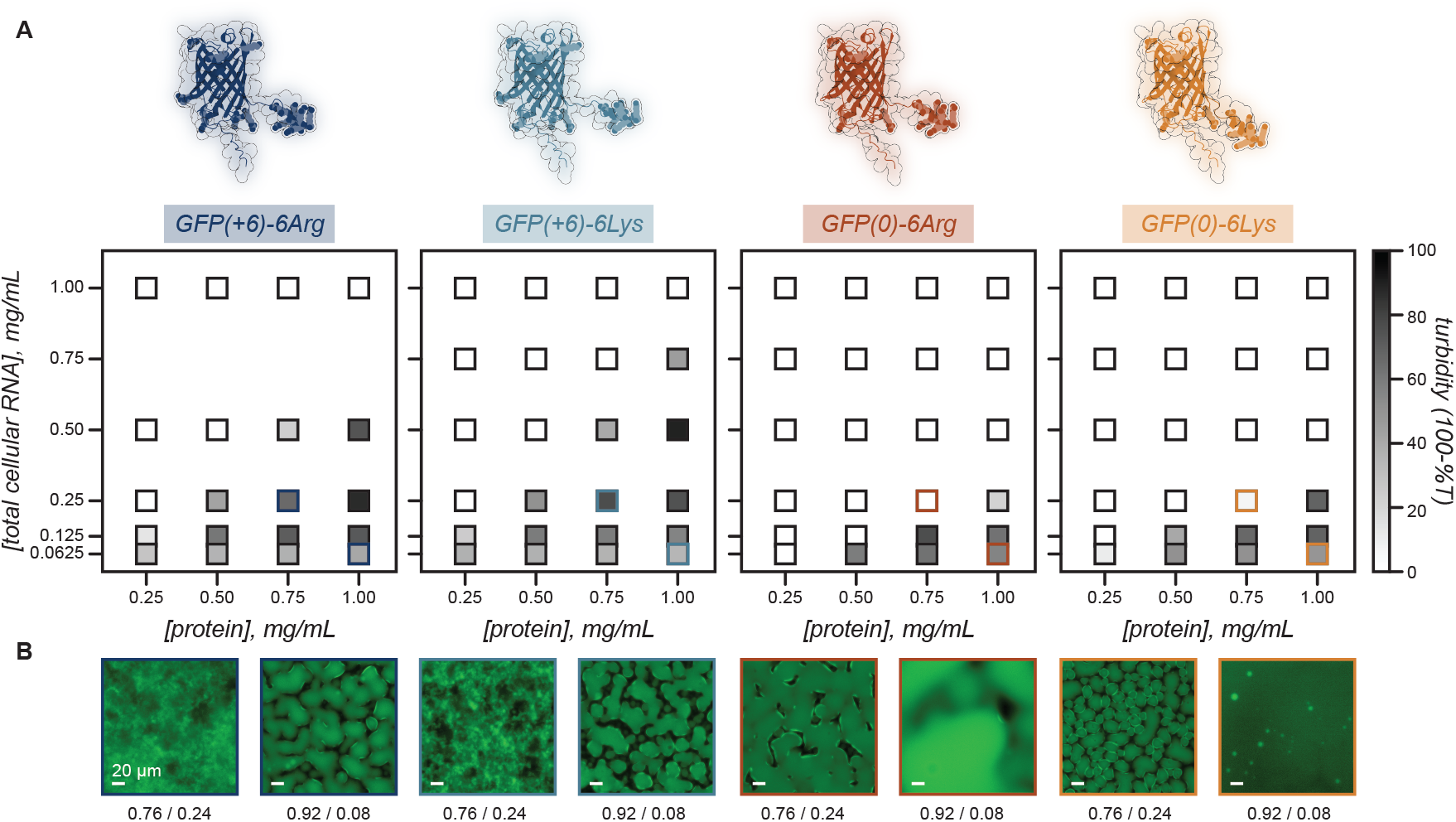
*In vitro* phase behavior of engineered GFP variants. (A, top) Design of tagged GFP(0) and GFP(+6) to assess the impact of arginine and lysine on the phase behavior with tcRNA. Protein structures as predicted by AlphaFold with point mutations to the surface of wild type sfGFP creating a net charge of 0 and +6 in GFP(0) and GFP(+6) variants, respectively. 6 Arg and 6 Lys tags were appended onto the C terminus of GFP(0) and GFP(+6). (bottom) Two dimensional phase diagrams of GFP(+6)-6Arg (dark blue), GFP(+6)-6Lys (light blue), GFP(0)-6Arg (dark orange) and GFP(0)-6Lys (light orange) with tcRNA from torula yeast. (B) Representative microscopy images of select mixing ratios from (A). Scale bar is 20 µm.

### Side chain composition controls phase behavior and salt resistance

We next wanted to confirm that the engineered proteins have the ability to undergo complex coacervation under physiological conditions. We individually mixed the four tagged proteins *in vitro* with total cellular RNA from torula yeast (tcRNA) at varying protein to RNA ratios to map out a phase diagram. Phase separation was observed in all constructs as monitored by solution turbidity^41^ and fluorescence microscopy (Fig 1A, B). The relative phase boundaries of these four proteins followed our hypothesis, with the GFP(+6) and arginine tagged proteins both exhibiting an expanded phase boundary when compared to GFP(0) and lysine tagged proteins, respectively. Both GFP(0) variants showed an increase in their C_sat_ values relative to GFP(+6) tagged proteins, indicating a higher protein concentration required for phase separation, likely due to the decreased electrostatic interactions relative to the more cationic GFP(+6) variants, which required lower protein concentrations to phase separate.

In these initial turbidity experiments, GFP(+6)-6Lys unexpectedly showed higher turbidity than the 6Arg variant at many conditions, despite prior evidence that arginine promotes phase separation relative to lysine^34,43^. By fluorescence microscopy, we observed that GFP(+6)-6Arg condensates were the only ones to form amorphous precipitates^44^ at the peak mixing ratios (Fig 1B). Liquid condensates scatter more than solid condensates, which can result in higher turbidity values that are not directly correlated to the extent of phase separation or degree of protein partitioning in the dense phase^44^. The impact of arginine versus lysine residues was further probed by interrogating the effect of NaCl, which disrupts electrostatic interactions, on coacervate formation (Fig 2A). At the critical concentration determined in the initial phase diagrams (1 mg/mL protein and 0.5 mg/mL tcRNA), GFP(+6)-6Arg exhibited increased turbidity in the presence of a small amount of NaCl (25 mM), than in the absence of NaCl (Fig 2A). The small disruption of the electrostatic interactions mediating protein:RNA complexation is sufficient to promote coacervates with more liquid-like properties that scatter more light, therefore leading to higher turbidity. Fluorescence microscopy confirmed the formation of liquid coacervates in the presence of NaCl for GFP(+6)-6Arg, as well as for all other arginine and lysine protein variants (SI Fig 4). Although increasing NaCl concentration was inversely correlated with turbidity values in all constructs, GFP(+6)-6Lys, GFP(0)-6Arg, and GFP(0)-6Lys did not show any liquid-to-solid or solid-to-liquid transitions (Fig 2A, SI Figure 2). Otherwise, only modest increases in the salt resistance of arginine-containing GFP variants was observed relative to the lysine tagged counterparts.

**Figure 2:**
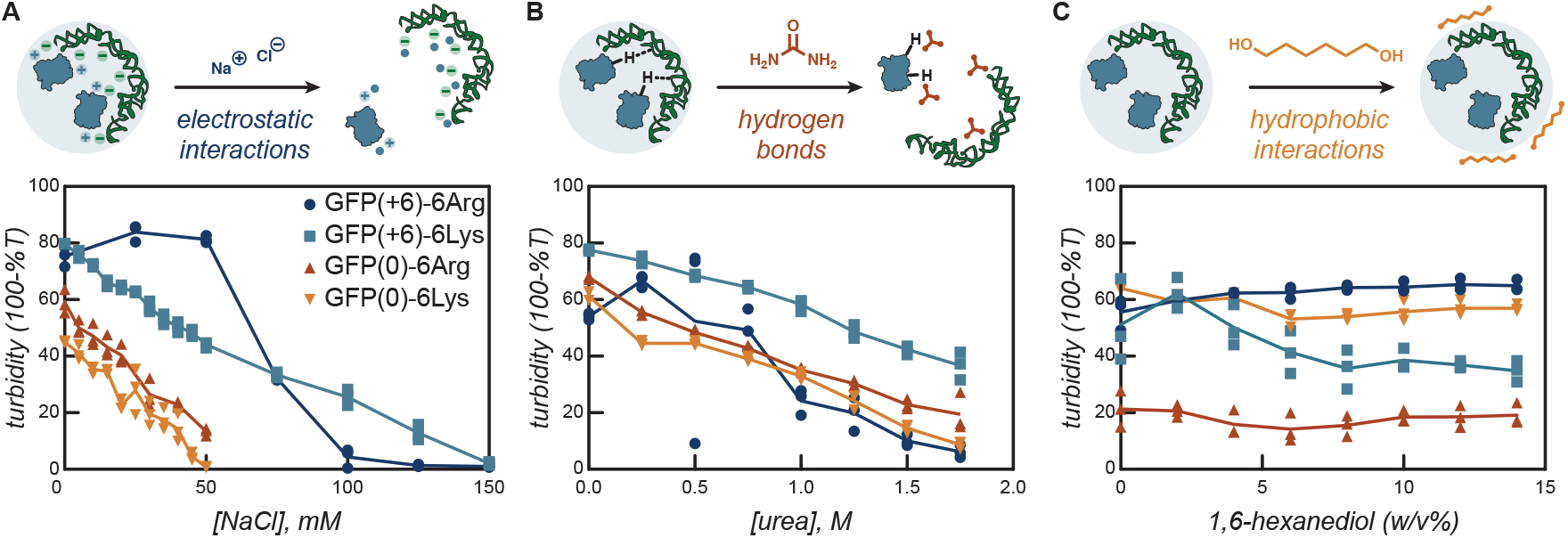
The impact of non-covalent interactions on the *in vitro* condensate formation of arginine and lysine tagged proteins with RNA. (A) Increasing NaCl concentration disrupts electrostatic interactions, decreasing condensate formation for all four protein variants. Proteins with a higher net charge (+12) show increase resistance to condensate dissolution as a function of salt concentration. There are only slight differences between Arg and Lys variants with a lower net charge of (+6). (B) Increasing concentrations of urea, which disrupts hydrogen bonding, shows equivalent impacts to all constructs. (C) Disruption of hydrophobic interactions via addition of 1,6-hexanediol shows no impact on turbidity values. Condensates in all experiments were prepared in 10 mM tris buffer at pH 7.4, and GFP(+6)-6Arg and GFP(+6)-6Lys (1.0 mg/mL) were mixed with tcRNA (0.5 mg/mL), while GFP(0)-6Arg and GFP(0)-6Lys (0.75 mg/mL) were mixed with tcRNA (0.125 mg/mL).

In addition to probing potential differences in electrostatic interactions, we also probed the role of hydrogen bonds and hydrophobic interactions in observed differences in coacervates formed with arginine and lysine tagged proteins. In a similar assay using urea, which disrupts hydrogen bonding^45^, identical behavior was observed between all four tagged constructs (Fig 2B). All four of the protein variants showed a consistent decrease in turbidity, with limited differences based on the protein net charge or identity of the cationic amino acid. Hydrophobic interactions were shown to be an insignificant contributor to the observed phase transitions via 1,6-hexanediol titration assays (Fig 2C)^45^. The addition of increasing volumes of this hydrophobicity modulator resulted in no changes in the solution turbidity, indicating that the formation of the condensed phase was not reliant on hydrophobic interactions. These combined results suggest that electrostatics are the non-covalent interaction that is most affected by alterations to protein cationic identity, and is also responsible for the observed hierarchy in salt stability and turbidity of all four tagged protein variants.

### *In vivo* characterization of phase behavior

After probing the *in vitro* characteristics of these synthetic condensates comprised of engineered protein and total cellular RNA, we next wanted to interrogate if the properties observed *in vitro* were also observed in the crowded cellular milieu. To do this, we utilized a BL21 strain of *E. coli* (NiCo21), as a simple prokaryotic model organism to directly probe the specific impact of side chain identity and net charge on phase separation.

Using fluorescence microscopy, we observed condensate formation in *E. coli* for all arginine and lysine tagged constructs with some distinct differences between the protein variants immediately apparent (Fig 3A, SI Figure 3). Using quantitative image analysis, we can approximate the ratio of protein in the synthetic condensates to that in the cytoplasm. We observe a higher condensate-to-cytoplasm ratio in cells expressing proteins with higher net charge or with arginine tags, with GFP(+6)-6Arg showing the highest condensate partitioning (Fig 3B). For example, for proteins with a net charge of +6, GFP(0)-6Arg shows increased partitioning in the synthetic condensates relative to GFP(0)-6Lys, indicating the enhanced interactions available for arginine. This arginine tagged variant also shows increased partitioning relative to the isotropically supercharged GFP(+6). Using this same approach, we can quantify the fraction of cells with condensates at a range of effective partition coefficients (SI Figure 4). All of the tagged protein variants had a high fraction of condensate containing cells (∼ 75-100%), as defined by a fluorescence ratio of at least 1.2 in the condensate relative to the cytoplasm, where the cytoplasm intensity was approximated by an average of 9 central pixels within the cell. However, if this threshold ratio for identifying condensates is increased even slightly to 1.4 or 2.0, we observe that the fraction of cells with condensates is higher for +12 charged variants (∼ 80-90%) than for +6 charged variants (∼ 10-80%). For the GFP(0) variants, we observe that the arginine tagged GFP consistently shows a higher fraction of condensate containing cells. These results show that the *in vivo* behavior of condensate formation largely matches the *in vitro* behavior observed, and that *E. coli* cells can be used as a model organism for the characterization of protein phase behavior^46^. Additionally, these findings support prior work from our group showing that cationic polypeptide tags appended to globular proteins such as GFP result in observable condensate formation that has charge-dependent behavior^47,48^. However, here we show that the identity of the cationic amino acid also plays a key role in dictating observed *in vivo* phase behavior.

**Figure 3:**
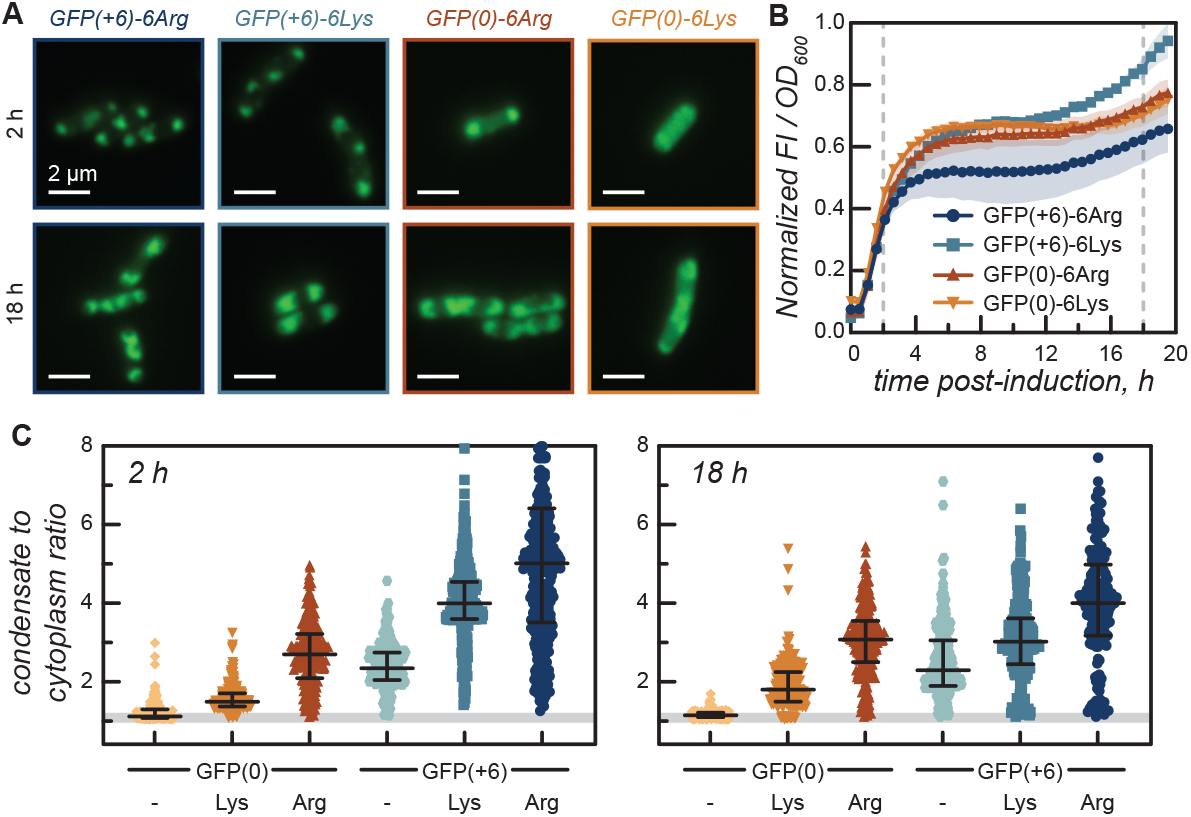
Condensate formation in *E. coli* cells. (A) Representative microscopy images of cells expressing arginine and lysine variants of GFP. (B) Protein expression was monitored by the FI488/OD600. The plot shows the relative protein expression from 0 to 20 h post induction for GFP(+6)-6Arg (dark blue), GFP(+6)-6Lys (light blue), GFP(0)-6Arg (dark orange) and GFP(0)-6Lys (light orange). Grey dashed lines indicate time points monitored by fluorescence microscopy. (C) Condensate to cytoplasm ratio analysis of individual cells across three biological replicates. The formation of protein condensates in *E. coli* was evaluated at 2 h and 18 h post induction of protein expression. Images were collected on a 100X oil 1.40 NA UPlanSApo objective, pre-processed in FIJI using the microbeJ plugin, and analyzed with Matlab.

It was also revealed that the condensate-to-cytoplasm ratio decreased slightly for all the GFP(+6) variants at long time points during the stationary phase (18 h) compared to an early time point during the exponential growth phase (2 h). Conversely, this ratio slightly increased for all GFP(0) variants (Fig 3B and SI Figure 4). We hypothesize that this is in large part due to the sequestration of essential mRNA and rRNA into phase separated condensates, with the GFP(+6) variants more effectively sequestering these anionic RNA biopolymers (Fig 5B-D)^49–52^. The sequestration of these essential biomolecules from the rest of the endogenous cellular machinery results in some cytotoxicity for strains that have strong condensate formation, as shown by decreased growth on agar plates (SI Figure 5). We propose that this cytotoxicity and resulting changes in the intracellular environment, likely explains the decrease in condensate-to-cytoplasm ratio of GFP(+6) tagged variants from 2 h to 18 h post-induction, while the other strains had a slight increase in this ratio as the protein concentration approximately doubled from 2 to 18 h post induction. But, there was no clear correlation found between the intracellular protein concentration, as approximated by the fluorescence intensity normalized by the optical density (FI/OD600), and the condensate-to-cytoplasm ratio for the four tagged variants (Fig 3C). Condensate-to-cytoplasm ratio was generally observed to coincide with growth and increased protein expression, but variations in protein expression levels alone does not play a noteworthy role in the hierarchical charge and side chain identity characteristics we observed.

### Cationic side chain identity influences *in vivo* protein mobility

Next, we wanted to understand how the observed differences in condensate formation in *E. coli* and protein partitioning in the condensed phase impacts protein mobility within these condensates. As previously mentioned, there are no statistically significant differences in protein expression levels (Fig 3B), suggesting that any differences observed regarding protein mobility are a result of the changes in amino acid identity influencing the interaction strength of condensate-bound molecules, and not a concentration-dependent behavior. To assess protein mobility in the condensates, we performed fluorescence recovery after photobleaching (FRAP) experiments, wherein approximately one condensate per cell was photobleached and then recovery at that same location was monitored. Overall, we found that condensates driven by arginine interactions were less mobile than those driven by lysine interactions, in agreement with previous findings^31,34^. For example, we found that, on average, it took GFP(+6)-6Lys based coacervates 18 s to reach 50% fluorescence at 2 h post induction, whereas GFP(+6)-6Arg based coacervates recovered to 50% fluorescence in 59 s at the same time point; nearly 3.3 times longer due to the simple change in the side chain chemistry of arginine. At 18 h post induction, neither construct reached 50% recovery. However, it took GFP(+6)-6Arg a similar 3.25 times longer (60 s) to reach 25% recovery as compared to GFP(+6)-6Lys (18.5 s). These differences are also reflected in the *t*_1*/*2_ values for the exponential fits of the recovery curves, with GFP(+6)-6Arg showing significantly slower recovery at 2 h than GFP(+6)-6Lys. Critically, the difference between lysine and arginine was also apparent for proteins of different net charge, with GFP(+6)-6Lys recovering more quickly than GFP(0)-6Arg.

We were unable to compare GFP(0)-6Arg with GFP(0)-6Lys because the latter construct did not reliably form condensates in *E. coli* for FRAP; the rapid diffusion of free protein in the cytoplasm resulted in 100% photobleaching during the bleach phase. However GFP(0)-6Arg condensates did show similar rates of recovery as those formed from GFP(+6)-6Lys, with the slower recovery dynamics of the lower charge arginine variant supporting the increased interaction strength of arginine relative to lysine (Fig4C).

**Figure 4:**
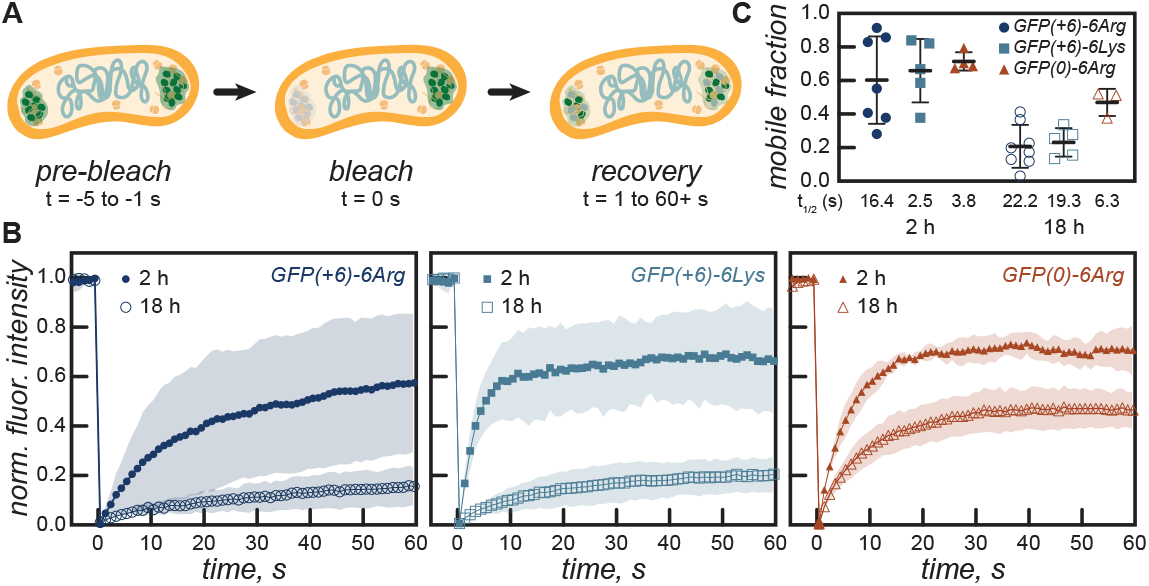
Dynamics of engineered protein condensates as determined by FRAP. (A) Schematic of FRAP in a model *E. coli* cell, showing pre-bleach, laser bleaching, and the recovery phases. (B) FRAP recovery curves of GFP(+6)-6Arg, GFP(+6)-6Lys, and GFP(0)-6Arg. GFP(0)-6Lys did not form sufficient condensates in *E. coli* for FRAP measurements. (C) A plot of the mobile protein fraction for the GFP variants that formed condensates at 2 and 18 h, as well as average t_1/2_ values for fluorescence recovery.

In addition to the timescales for fluorescence recovery being impacted by both the protein net charge and the identity of the cationic amino acid, we also find that the mobile fraction within the condensate was also dependent on these features. Using the equation,

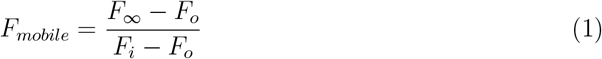

we find the mobile fraction of GFP(+6)-6Arg at 2 h post induction to be 60%, which is approximately 6% lower than GFP(+6)-6Lys at 2 h post induction (66%). The mobile fraction of GFP(+6)-6Arg drops to 21% at 18 h post induction, while GFP(+6)-6Lys drops to 23%, However, despite the lower valency relative to GFP(+6)-6Lys, the mobile fraction of GFP(0)-6Arg was only a little over 8% greater than that of GFP(+6)-6Lys at 2 h post induction. The mobile fraction of all constructs decreased from 2 h to 18 h, as expected, but GFP(0)-6Arg exhibited the least significant drop in the mobile fraction; dropping from 71% at 2 h to 47% at 18 h.

### Interactions of nucleic acids with condensates *in vivo*

After analysis of *in vivo* protein partitioning in the condensate for the arginine and lysine GFP variants, it still remained unclear if the *in vivo* condensates observed were a product of heterotypic complex coacervation with nucleic acids. Our *in vitro* experiments utilized RNA to complex with the tagged protein constructs, so we wanted to determine if these protein-RNA contacts were also the dominant interaction mediating condensate formation in *E. coli*.

To visualize RNAs in *E. coli* we use the mPepper aptamer^50–52^ to fluorescently tag RNA (Fig 5A). We included four repeats of the mPepper aptamer sequence at the 3’ end of mRNA encoding for the engineered protein variants. Staining with the HBC620 ligand, allowed us to observe mRNA colocalization with all condensates. Additionally, simultaneous staining with DAPI, a DNA selective probe, showed the exclusion of DNA from all condensates (Fig 5B, C). As condensate formation was less pronounced for GFP(0)-6Arg and GFP(0)-6Lys variants, the exclusion of the DNA from these condensates was slightly less clear than for the GFP(+6) tagged variants. Cells without condensate formation exhibited homogeneous intensity profiles for the GFP, mPepper and DAPI signals.

**Figure 5:**
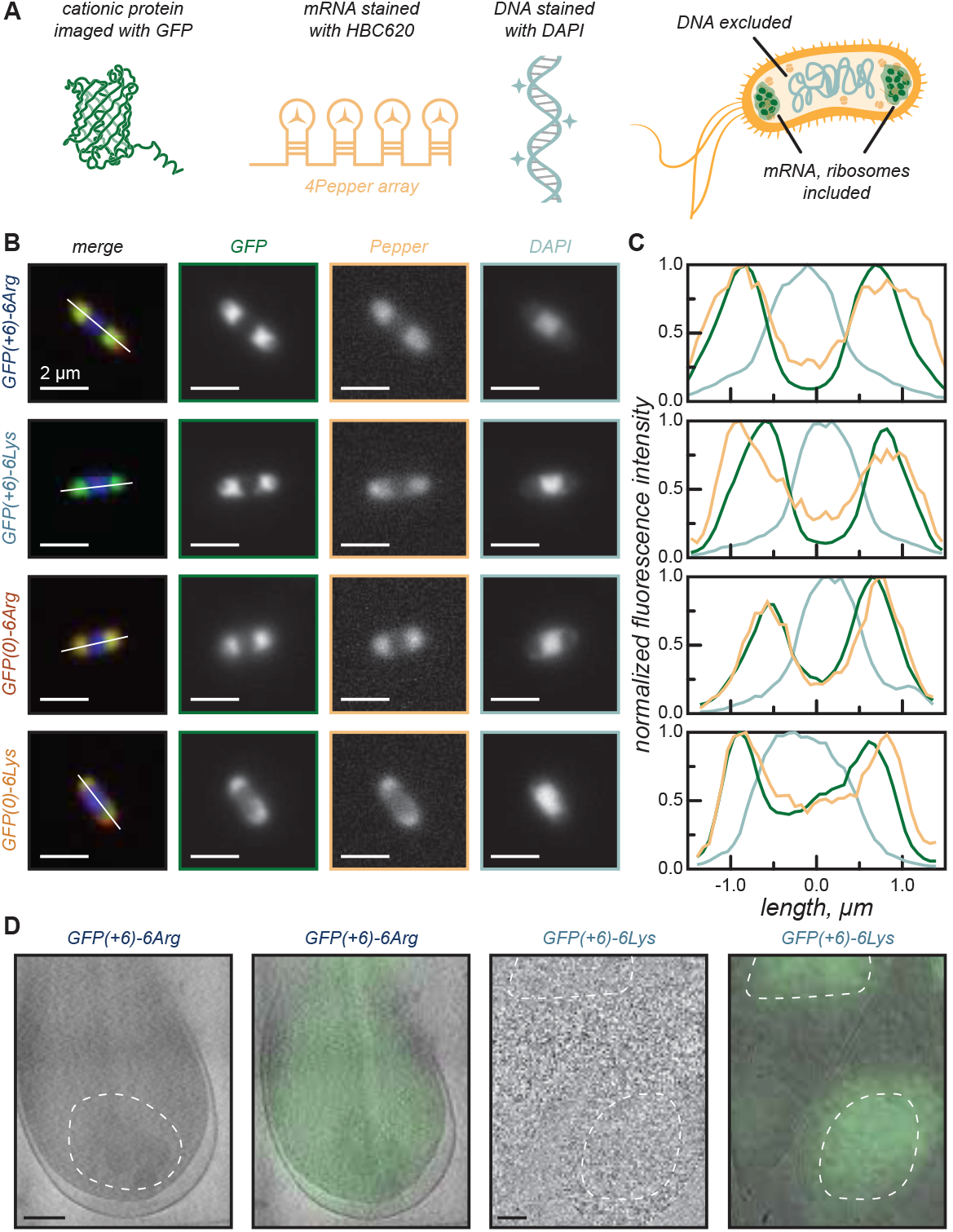
The distribution of fluorescently tagged nucleic acids in *E. coli* with engineered protein condensates. (A) Schematic showing how different proteins and nucleic acids were visualized and how they localize in *E. coli* with condensates. (B) Representative fluorescent microscopy images of GFP(+6)-6Arg, GFP(+6)-6Lys, GFP(0)-6Arg, and GFP(0)-6Lys showing localization of GFP and mPepper in the condensates, as well as DAPI stained DNA exclusion from the condensates. (C) Normalized fluorescence intensity plot profiles measured on cells with two condensates, with one at each pole. Orange lines represent mPepper, light blue lines represent DAPI, and green lines represent GFP fluorescence. (D) Cryo-CLEM overlays of GFP(+6)-6Arg and GFP(+6)-6Lys, showing that the region of high electron density in the cryoEM slice is co-localized with the GFP fluorescence signal.

In addition to visualizing mRNA localization, we also used high resolution cryo-correlative light and electron microscopy (CLEM) imaging to better visualize condensates and ribosomes within the cells. GFP(+6) samples that reliably formed condensates were first imaged by confocal microscopy and then subsequently imaged with high-resolution cryo-electron tomography. We observe that the fluorescence signal is co-localized with distinct regions containing amorphous-like density. Initial tomogram reconstructions showed condensates as amorphous regions of high electron density, potentially indicating colocalization of ribosomes within condensates (Fig 5D). However, tomography methods utilizing denoising methods, as done by IsoNet, resulted in the observation of a void region for GFP(+6)-6Lys condensates instead (Fig 5D), Supplementary Fig 7), indicating that these regions were rather disordered.

## Conclusions

The current understanding of the molecular determinants for biomolecular condensate formation is best described as a rapidly expanding encyclopedia. Valency^41^, molecular weight^34^, post translational modifications^33^, degree of disorder^20^, small molecules^53^, general stress^54^, and local buffer conditions^55^ have all been shown to play a role in influencing the formation and dynamics of condensates. The recent appreciation of biomolecule condensation, often described as liquid-liquid phase separation, as an important phenomena in many biological processes^2^, efforts have focused on understanding the laws governing when, where, why, and how it occurs. A further grasp of the molecular mechanisms governing phase separation has already lead to new approaches for treating a range of diseases from cancer to viral infections and previously gridlocked neurodegenerative conditions^16–18^. Understanding these principles at a molecular level allows for the ground-up engineering of synthetic systems, modeled from biomolecular condensates, which have applications spanning from drug delivery systems, to protein purification^27,40^. Here, we sought to evaluate how cationic amino acid composition regulates protein dynamics and phase behavior in condensates in complexation with RNA, using the two positively charged amino acids at physiological pH, lysine and arginine. We showed important variation in the phase behavior driven by the two amino acids. Using total cellular RNA from torula yeast to mimic a complete eukaryotic transcriptome, we observed the formation of arginine-based condensates that show a greater resistance to salt as compared to lysine-based coacervates. While in low salt conditions the phase boundaries of arginine-tagged and lysine-tagged protein variants were similar, the greatest difference in condensate formation was observed as we tuned the strength of electrostatic interactions by titrating the NaCl concentration. In the presence of salt, we observed an expanded two phase region in both the GFP(0) and GFP(+6) arginine constructs compared to the lysine constructs. This shows the stronger electrostatic interactions between arginine and tcRNA, likely due to the *π*-interactions arginine is able to engage in (*π* − *π*, cation-*π*). Using additional titration assays, we did not observe that any other non-covalent interaction significantly contributed to the observed differences in phase separation.

We also demonstrated in *E. coli* that arginine tagged constructs showed increased condensate formation. We showed that arginine protein variants had higher partitioning in the condensed phase, relative to the lysine containing variants. Notably, we observed the colocalization of the GFP-tagged proteins with RNA inside *E. coli* condensates, while excluding genomic DNA, suggesting the preferred complexation partner is RNA, as has been observed in previous reports^42^. Furthermore, the protein mobility within these condensates was also dependent on the arginine content, with arginine variants consistently showing slower recovery. These results show how varying cationic side chain identity significantly impacts condensate behavior with near equal importance to the overall charge of the protein. Importantly, this provides an additional design parameter to tune when designing synthetic biomolecular condensates.

## Acknowledgments

A.K.K. and A.C.O acknowledge the National s of Health (NIH) for funding this work under an award from the National Institute of General Medical Sciences (NIGMS, R35GM138378). Additionally, these studies used the Confocal and Specialized Microscopy Shared Resource of the Herbert Irving Comprehensive Cancer Center at Columbia University, funded in part through the NIH/NCI Cancer Center Support Grant P30CA013696. Cryo-ET experiments were supported by the Simons Electron Microscopy Center and the National Resource for Automated Molecular Microscopy located at the New York Structural Biology Center, supported by grants from the Simons Foundation (SF349247) and the NIH NIGMS (GM103310).

## Conflict of Interest

The authors declare that there are no conflicts of interest associated with this manuscript.

## Methods

### Cloning of GFP(0) and GFP(+6) tagged Arg/Lys constructs

All primers for cloning were ordered from Integrated DNA Technologies (IDT). All template DNA for cloning experiments were ampicillin resistant and located on a pETDuet vector carrying either a 6xHis-GFP(0) or 6xHis-GFP(+6) gene inserted after the *NcoI* site. Primers used to PCR amplify DNA for HiFi assembly were created with the guidance of NEBuilder (New England Biolabs, NEB). PCR products were produced using Phusion high fidelity DNA polymerase (Thermo Scientific), resolved on agarose gels in TAE buffer, and purified from the gel using the QIAquick Gel Extraction Kit (Qiagen). Purified PCR products were assembled into plasmids using the NEB HiFi DNA assembly kit following the manufacturer’s protocol, and subsequently transformed into NEB5*α* competent cells using the high efficiency transformation protocol from the manufacturer (NEB). Transformed plasmids were sequence verified using Sanger sequencing (Genewiz). Sequence verified plasmids were then transformed into NiCo21 (DE3) competent cells using the high efficiency transformation protocol (NEB C2529).

### Protein Expression

Overnight cultures (50 mL) of NiCo21 (DE3) cells harboring plasmids for the expression of GFP variants were grown for approximately 15 h in LB media supplemented with ampicillin (100 µg/mL) in Erlenmeyer flasks from a colony selected from an LB agar plate supplemented with ampicillin. Cultures were grown at 37 °C, with shaking at 225 rpm. After the initial growth overnight, the culture was used to inoculate a 1 L culture of LB media supplemented with ampicillin (100 µg/mL). The cells were incubated at 37 °C with shaking at 225 rpm until an optical density (OD_600_) of 0.7-0.9 was reached. At that point, isopropyl ß-D-1-thiogalactopyranoside (IPTG) was added to a final concentration of 1 mM and the cultures were grown at 37 °C, with shaking at 225 rpm.

### Protein Purification

Cells were harvested from cultures 16-18 h post induction via centrifugation (4000 rpm for 10-20 min). The supernatant was discarded and pelleted cells were resuspended in 15-30 mL lysis buffer (50 mM NaH_2_PO_4_, 300 mM NaCl, pH 8.0) per L of culture. Cells were sonicated three times at 60% amplitude with a 2s on, 4s off cycle for 5 min (Fisher Scientific FB505). Sonication was performed in 50 mL tubes submerged in ice water. Cell debris was subsequently removed by centrifugation (13,000 rpm for 30 min). The supernatant was carefully decanted into a new 50 mL tube and all pelleted debris was discarded.

This soluble protein fraction was purified using Ni-NTA metal affinity chromatography (HisPur Ni-NTA Resin (Thermo Scientific, 88223)) following a general protocol from the manufacturer. Briefly, 15-30 mL of Ni-NTA resin in a 50:50 slurry with 20% ethanol was used per L of culture. After equilibrating the beads in lysis buffer, the beads were incubated at room temperature for 10 min with the clarified supernatant. The flow through was collected and the resin was then rinsed 3x times with 70 mL lysis buffer. The beads were washed 5 times with 50 mL wash buffer (lysis buffer with 30 mM imidazole), and finally protein was eluted from the beads with elution buffer (lysis buffer with 250 mM Imidazole) until the eluate ran clear.

The flow through, wash, and elution fractions were analyzed by SDS-PAGE. Pure fractions were combined and concentrated by centrifugal ultrafiltration with a 10 kDa molecular weight cutoff (MWCO) filter. Following concentration, protein solutions were dialyzed against a 10 mM tris buffer at pH 7.4. After dialysis, the protein concentration was adjusted to 1 mg mL^-1^ for further experiments. The identity of the purified proteins was confirmed by MALDI-TOF mass spectrometry. Samples were prepared for MALDI-TOF analysis by performing a 10-fold dilution of the 1 mg mL^-1^ stock protein solution into MilliQ water. A saturated solution of sinapinic acid (10 mg mL^-1^) in a 7:3 mixture of water and acetonitrile was used as the matrix solution. Final samples were prepared with 70% matrix solution and 30% protein solution prior to spotting 1 *µ*L on a stainless steel plate.

### *E. coli* cell growth for fluorescence and electron microscopy imaging

A single colony of NiCo21 (DE3) cells (NEB) with a plasmid of interest was selected from an LB-agar plate, inoculated in 8 mL of LB media supplemented with ampicillin, and allowed to grow for 17 h at 37 °C with shaking at 225 rpm. The next day the cells were back diluted into 50 mL of fresh LB media containing ampicillin in a 125 mL Erlenmeyer flask to a final OD_600_ of 0.2. The cells were then grown at 37 °C, with shaking at 225 rpm, until they reached an OD_600_ of 0.8-1.0, at which point protein production was induced by supplementing IPTG to a final concentration of 1 mM. Cells were grown until time points for microscopy were reached.

### Fluorescence Microscopy of *E. coli*

*E. coli* cells were imaged on an agarose pad. 50 µL of a 1.5 w/v% agarose solution in water was pipetted on a 25×75 mm microscopy slide (Globe Scientific 1301). A coverslip (No. 1.5 18×18 mm, Thermo Scientific) was placed on the agarose pad as it cooled so as to leave a level surface for imaging. Once solidified, the cover slip was gently removed and 1-2 µL of cell culture was pipetted on the pad, the cover slip was re-applied, and the cover slip was sealed with a thin coat of clear nail polish (Electron Microscopy Sciences 72180) and imaged on an EVOS FL Auto 2 inverted fluorescence microscope (Invitrogen). Cells were imaged using a 100X oil 1.40 NA UPlanSApo objective (Olympus) on GFP (*λ*_ex_ = 470-522 nm; *λ*_em_ = 525-550 nm; EVOS GFP light cube) and brightfield channels. Cells were imaged on agarose pads at 2 and 18 h post-induction. Between 7-15 images were acquired for each strain at each time point in order to image and analyze a minimum of 45 cells per replicate.

### Image analysis

After collecting fluorescence images, the TIFF files were imported to FiJi (ImageJ). Stacks of the GFP channel images were created for each cell strain at each time point. Background subtraction with a rolling ball radius of 100 pixels was performed to correct for background signal. The plugin MicrobeJ (Version 5.13m (16)) was then used to identify cells and measure intracellular fluorescence. Images were thresholded using the Li method and constraints were placed on cell area (300-4500 p^2^), length (≥ 15 p), width (≤ 28 p), circularity (≥ 0.25), and angularity (≤ 0.55). After appropriate thresholding values were set, the dataset of identified cells was loaded into MATLAB and a custom script was used to identify the fraction of cells that contained condensates as a function of the ratio of condensate to cytoplasm^47^.

### DAPI and HBC620 staining

DAPI (Thermo-Fisher 62248, *λ*_ex_= 360 nm *λ*_em_= 460 nm) and HBC620 (FD Biotech Cat No. H16201, *λ*_ex_= 570 nm *λ*_em_= 620 nm) stains were used for the fluorescent labeling of intracellular nucleic acids. Cells for microscopy were inoculated, grown overnight, and induced with IPTG as described above. Before cells were placed on the agar pad for imaging, an aliquot of 250 *µ*L was placed in an eppendorf tube and centrifuged for 5 min at 13,000 rpm to pellet the cells. The growth medium was decanted by pipetting and 250 *µ*L of fresh phosphate buffered saline (PBS: 10 mM NaH_2_PO_4_, 1.8 mM KH_2_PO_4_, 2.7 mM KCl 140 mM NaCl, pH 7.4) was used to resuspend the pelleted cells by gentle pipetting. At this point, 0.5 µL of a 5 mg mL^-1^ DAPI solution in dimethylformamide (DMF) was added to the resuspended cells and mixed by pipetting. For HBC620, a PBS solution containing 2 *µ*M HBC620 was created, and the pelleted cells were resuspended in 250 µl of this solution by vortexing. The dye and cells are allowed to incubate at room temperature in complete darkness for 10 min before being imaged as above. For cells incubated with both dyes, the pellet was resuspended in PBS buffer containing 2 *µ*M HBC620 and then supplemented with DAPI

### Fluorescent recovery after Photobleaching (FRAP)

FRAP experiments were conducted at the Confocal and Specialized Microscopy Shared Resource of the Herbert Irving Comprehensive Cancer Center at Columbia University Medical Center. Images were collected on a Nikon A1 RMP/ Ti confocal microscope with a 100X 1.45 NA Plan-Apochromat oil immersion objective on GFP (*λ*_ex_= 488 nm; *λ*_em_= 500-550 nm) and brightfield (TD) channels. A circular area 0.2 µm in diameter was bleached with a 487 nm laser at 75% power for 0.0625 s. Post-bleach frames were captured using illumination with a 487 nm laser at 0.5% power. Recovery curves were fit to a single exponential recovery function using easyFRAP and averaged over 4-8 cells. Replicates with a gap ratio and/or bleach depth below 0.6 were excluded from analysis because of excessive or insufficient bleaching. The mobile fraction and t_1/2_ values were obtained from nonlinear fit analysis performed in Graphpad Prism 10.

### LB-Agar Spotting Assay

LB-agar plates were poured into Nunc™ OmniTray™ Single-Well Plates (ThermoFisher 165218). Control plates were supplemented with ampicillin and experimental plates were supplemented with ampicillin and IPTG (1 mM). Strains were grown concurrently following the same protocol as detailed above until all constructs reached an OD_600_ ≥ 0.9. At that point all constructs were back-diluted to an OD_600_=0.9 exactly in the wells of the first column of a 96-well plate, the cultures were then serially diluted 1:2 in LB media supplemented with ampicillin for a total of 9 times. A 96-pin microplate replicator (VP Scientific 407) was dipped into the 96-well plate for 5 s, and then pressed directly onto the agar plate for 5 s without disturbing the agar. After repeating the spotting on control plates, the plates were grown at 37 °C for 18 h. Plates were then imaged using an Epson perfection V850 scanner and analyzed for intensity of the resulting colonies using FiJi.

### Sample Preparation and Cryo-FIB Milling

Sample preparation and cryo-focused-ion beam (cryo-FIB) milling for cryo-ET studies was performed using the Waffle method^56,57^, at the New York Structural Biology Center (NYSBC).

∼

Briefly, cell cultures were centrifuged at 500 × g for 10 min, and the supernatant was removed. The cell pellet was then resuspended in 100 *µ*L of LB media containing 10% glycerol. 3 *µ*L of this cell suspension was then applied onto Glow-discharged (for 40 s at 1 mA with the grid bar side placed facing up) Quantifoil R2/1 grids with 200 copper meshes that were coated with an extra film of carbon. The waffle grids were assembled in a platinum planchette and were subjected to high-pressure freezing with an Wohlwend HPF compact 01 high-pressure freezer. The sample was loaded on an Aquilos2 system for cryo-FIB milling on to a shuttle with a 45^*°*^ pre-tilt and were sputter coated for 10 s. An organometallic platinum GIS layer was deposited with a gas injection system for ∼ 3 min. AutoTEM (v.2.0-2.3, ThermoFisher Scientific) was used for all subsequent steps. Precuts were milled following the waffle method protocol,^56,57^ with a beam current of 10 nA at 30 kV. The eucentric position and milling position for each lamella site was defined and the lamella was subsequently cleaned by gradually reducing the milling angles. At a final 20^*°*^milling angle, a notch pattern,^56,57^ was milled with a beam current of 0.3 nA for ∼ 2 min. AutoTEM was used to automate the milling at each lamella site, aiming for a final thickness of 180-300 nm at a 20^0^ milling angle.

### Cryo-ET Data Collection, Processing, and Correlative-Light Microscopy

Fluorescent images for GFP(+6)-6Arg, GFP(+6)-6Lys, and GFP(0) (control) lamella were collected on a Zeiss LSM 900 with Airyscan 2 LSM980 confocal microscope, before cryo-ET data collection as the radiation damage removed the fluorescent signal from the lamella. 100x magnification Z-stacks were collected on each lamella, with the excitation wavelength set to 488 nm and emission wavelength set to 509 nm, and were aligned using wavelets set to highest in the Zeiss Zen software. Data collection for all lamella was performed on a Titan Krios microscope (ThermoFisher Scientific) equipped with a field emission gun, a GIF quantum LS post column energy filter (Gatan) and a K3 summit electron detector (Gatan). The electron microscope was operated at 300 kV in nanoprobe mode at a magnification from 19500 to 22000k (nominal pixel sizes from 3.55-4.5 ^Å^ at the specimen level). Cryo-ET tilt series were collected using a dose-symmetric scheme^58^ with a tilt-range from − 45^*°*^ to +45^*°*^ degrees after tilting stage to 20^*°*^ to account for the milling angle of the lamella. The tilt-series were collected at a defocus of -4 *µ*m to -8 *µ*m, in 3^*°*^ tilts for the arginine and lysine tagged constructs and 2^*°*^ tilts for the GFP(0) control construct, with a total dose of 135 e/^Å2^ with SerialEM^59^.

The raw tilt-series were motion corrected in Warp (v1.09)^60^. The tilt-series were exported from Warp and subsequently aligned and reconstructed with the software package AreTomo^61^. Missing-wedge correction and denoising of the tomograms was performed using IsoNet^62^. Medium magnification montages were blended using the blendmont command in the software Imod^63^ with default settings. For the purpose of overlaying the GFP signals on the blended montages and tomograms, Adobe Photoshop was used. Blended montages had the contrast increased, but were otherwise left unaltered. The GFP images were then added as additional layers, had their brightness increased, and their fill reduced to around 20%. Using the Scale, Rotate and Skew tools, the GFP images were manually aligned with the blended montages. For each tomogram, 3dmod was used to find the z-height which most closely matched the corresponding area on the blended montage. This image was added as a new layer, then resized and rotated to match the area. The GFP signal that overlapped with the tomogram was copied, rotated and resized back to the tomogram’s original configuration, then overlaid on top of the tomogram.

## Notes

### Competing Interest Statement

The authors have declared no competing interest.

