## Supplemental Information for "Cationic amino acid identity and net charge influence condensate properties in *E. coli*"

Kidane *et al.*

### Section 1: Supplemental Experimental Details

Table 1. DNA Sequences

| | |
|---|---|
| GFP(0) | ATGGGTCATCACCACCACCATCACGGTGGCGCTAGCAAAGGTGAAC<br>GTCTGTTTACTGGTGTAGTACCGATCTTAGTGGAATTAGACGGCGAC<br>GTGAACGGTCATAAATTTAGCGTGCGCGGCGAGGGCGAAGGTGACG<br>CTACCAATGGTAAATTGACCCTGAAGTTTATTTGCACAACAGGCAAA<br>TTACCCGTTCCGTGGCCCACCTTAGTGACCACCCTGACCTATGGCGTT<br>CAGTGCTTCAGTCGTTACCCTGATCATATGAAACAACACGATTTTTC<br>AAATCAGCCATGCCTGAAGGATATGTTCAAGAGCGTACAATCAGCTT<br>CAAGGACGATGGCACCTATAAAACGCGTGCGGAAGTGAAATTTGAA<br>GGCGACACATTAGTAAATCGTATCGAACTGAAAGGTCGTGACTTCAA<br>AGAAGACGGCAACATTTTAGGCCATAAACTGGAATATAACTTTAATT<br>CTCATAACGTGTATATTACGGCCGATAAACAGAAGAATGGTATCAAG<br>GCAAATTTCAAAATTCGCCATAACGTGGAGGACGGCAGCGTTCAATT<br>AGCGGATCATTATCAACAAAACACGCCGATTGGTGATGGGCCTGTAC<br>TGTTACCTCGCAACCACTACCTGAGCACCCAATCTGCCCTGAGCAAA<br>GATCCGAAAGAAAAACGCGATCACATGGTTCTGTTAGAATTCGTGAC<br>CGCTGCAGGCATTACGCACGGAATGGACGAACGCTACAAGTAA |
| GFP(0)-6Lys | ATGGGTCATCACCACCACCATCACGGTGGCGCTAGCAAAGGTGAAC<br>GTCTGTTTACTGGTGTAGTACCGATCTTAGTGGAATTAGACGGCGAC<br>GTGAACGGTCATAAATTTAGCGTGCGCGGCGAGGGCGAAGGTGACG<br>CTACCAATGGTAAATTGACCCTGAAGTTTATTTGCACAACAGGCAAA<br>TTACCCGTTCCGTGGCCCACCTTAGTGACCACCCTGACCTATGGCGTT<br>CAGTGCTTCAGTCGTTACCCTGATCATATGAAACAACACGATTTTTC<br>AAATCAGCCATGCCTGAAGGATATGTTCAAGAGCGTACAATCAGCTT<br>CAAGGACGATGGCACCTATAAAACGCGTGCGGAAGTGAAATTTGAA<br>GGCGACACATTAGTAAATCGTATCGAACTGAAAGGTCGTGACTTCAA<br>AGAAGACGGCAACATTTTAGGCCATAAACTGGAATATAACTTTAATT<br>CTCATAACGTGTATATTACGGCCGATAAACAGAAGAATGGTATCAAG<br>GCAAATTTCAAAATTCGCCATAACGTGGAGGACGGCAGCGTTCAATT<br>AGCGGATCATTATCAACAAAACACGCCGATTGGTGATGGGCCTGTAC<br>TGTTACCTCGCAACCACTACCTGAGCACCCAATCTGCCCTGAGCAAA<br>GATCCGAAAGAAAAACGCGATCACATGGTTCTGTTAGAATTCGTGAC<br>CGCTGCAGGCATTACGCACGGAATGGACGAACGCTACAAGAAAAAG<br>AAAAAGAAGAAATAA |
| R. Primer | 5’-TTTCTTCTTTTTCTTTTTCTTGTAGCGTTCGTCCATTCC-3’ |
| F. Primer | 5’- |

| | |
|---|---|
| | GAATGGACGAACGCTACAAGAAAAAGAAAAAGAAGAAATAATAATGAGGTACCCTCG-3’ |
| GFP(0)-6Arg | ATGGGTCATCACCACCACCATCACGGTGGCGCTAGCAAAGGTGAACGTCTGTTTACTGGTGTAGTACCGATCTTAGTGGAATTAGACGGCGACGTGAACGGTCATAAATTTAGCGTGCGCGGCGAGGGCGAAGGTGACGCTACCAATGGTAAATTGACCCTGAAGTTTATTTGCACAACAGGCAAATTACCCGTTCCGTGGCCCACCTTAGTGACCACCCTGACCTATGGCGTTCAGTGCTTCAGTCGTTACCCTGATCATATGAAACAACACGATTTTTCAAATCAGCCATGCCTGAAGGATATGTTCAAGAGCGTACAATCAGCTTCAAGGACGATGGCACCTATAAAACGCGTGCGGAAGTGAAATTTGAAGGCGACACATTAGTAAATCGTATCGAACTGAAAGGTCGTGACTTCAAAGAAGACGGCAACATTTTAGGCCATAAACTGGAATATAACTTTAATTCTCATAACGTGTATATTACGGCCGATAAACAGAAGAATGGTATCAAGGCAAATTTCAAAATTCGCCATAACGTGGAGGACGGCAGCGTTCAATTAGCGGATCATTATCAACAAAACACGCCGATTGGTGATGGGCCTGTACTGTTACCTCGCAACCACTACCTGAGCACCCAATCTGCCCTGAGCAAAGATCCGAAAGAAAAACGCGATCACATGGTTCTGTTAGAATTCGTGACCGCTGCAGGCATTACGCACGGAATGGACGAACGCTACAAGCGTCGCCGGCGCCGTCGGTAA |
| R. Primer | 5’-CCGACGGCGCCGGCGACGCTTGTAGCGTTCGTCCATTCC-3’ |
| F. Primer | 5’-GAATGGACGAACGCTACAAGCGTCGCCGGCGCCGTCGGTAATAATGAGGTACCCTCG-3’ |
| GFP(0)-6Lys 4Pepper | ATGGGTCATCACCACCACCATCACGGTGGCGCTAGCAAAGGTGAACGTCTGTTTACTGGTGTAGTACCGATCTTAGTGGAATTAGACGGCGACGTGAACGGTCATAAATTTAGCGTGCGCGGCGAGGGCGAAGGTGACGCTACCAATGGTAAATTGACCCTGAAGTTTATTTGCACAACAGGCAAATTACCCGTTCCGTGGCCCACCTTAGTGACCACCCTGACCTATGGCGTTCAGTGCTTCAGTCGTTACCCTGATCATATGAAACAACACGATTTTTCAAATCAGCCATGCCTGAAGGATATGTTCAAGAGCGTACAATCAGCTTCAAGGACGATGGCACCTATAAAACGCGTGCGGAAGTGAAATTTGAAGGCGACACATTAGTAAATCGTATCGAACTGAAAGGTCGTGACTTCAAAGAAGACGGCAACATTTTAGGCCATAAACTGGAATATAACTTTAATTCTCATAACGTGTATATTACGGCCGATAAACAGAAGAATGGTATCAAGGCAAATTTCAAAATTCGCCATAACGTGGAGGACGGCAGCGTTCAATTAGCGGATCATTATCAACAAAACACGCCGATTGGTGATGGGCCTGTACTGTTACCTCGCAACCACTACCTGAGCACCCAATCTGCCCTGAGCAAAGATCCGAAAGAAAAACGCGATCACATGGTTCTGTTAGAATTCGTGACCGCTGCAGGCATTACGCACGGAATGGACGAACGCTACAAGAAAAAGAAAAAGAAGAaataataatgaggtaccctcgagtctggtaaagaaaccgctgctgcgaaatgcccacg |

| | |
|---|---|
| | gaggatccccaatcgtggcgtgtcggcctctcccaatcgtggcgtgtcggcctctcccaatcgtggcgtgtcggcc<br>tctcccaatcgtggcgtgtcggcctct |
| GFP(0)-6Arg<br>4Pepper | ATGGGTCATCACCACCACCATCACGGTGGCGCTAGCAAAGGTGAAC<br>GTCTGTTTACTGGTGTAGTACCGATCTTAGTGGAATTAGACGGCGAC<br>GTGAACGGTCATAAATTTAGCGTGCGCGGCGAGGGCGAAGGTGACG<br>CTACCAATGGTAAATTGACCCTGAAGTTTATTTGCACAACAGGCAAA<br>TTACCCGTTCCGTGGCCCACCTTAGTGACCACCCTGACCTATGGCGTT<br>CAGTGCTTCAGTCGTTACCCTGATCATATGAAACAACACGATTTTTC<br>AAATCAGCCATGCCTGAAGGATATGTTCAAGAGCGTACAATCAGCTT<br>CAAGGACGATGGCACCTATAAAACGCGTGCGGAAGTGAAATTTGAA<br>GGCGACACATTAGTAAATCGTATCGAACTGAAAGGTCGTGACTTCAA<br>AGAAGACGGCAACATTTTAGGCCATAAACTGGAATATAACTTTAATT<br>CTCATAACGTGTATATTACGGCCGATAAACAGAAGAATGGTATCAAG<br>GCAAATTTCAAAATTCGCCATAACGTGGAGGACGGCAGCGTTCAATT<br>AGCGGATCATTATCAACAAAACACGCCGATTGGTGATGGGCCTGTAC<br>TGTTACCTCGCAACCACTACCTGAGCACCCAATCTGCCCTGAGCAAA<br>GATCCGAAAGAAAAACGCGATCACATGGTTCTGTTAGAATTCGTGAC<br>CGCTGCAGGCATTACGCACGGAATGGACGAACGCTACAAGCGTCGC<br>CGGCGCCGTCGGTaataatgaggtaccctcgagtctggtaaagaaaccgctgctgcgaaatgcccac<br>ggaggatccccaatcgtggcgtgtcggcctctcccaatcgtggcgtgtcggcctctcccaatcgtggcgtgtcggc<br>ctctcccaatcgtggcgtgtcggcctct |
| GFP(+6) | ATGGGTCATCACCACCACCATCACGGTGGCGCTAGCAAAGGTGAAC<br>GTCTGTTTACTGGTGTAGTACCGATCTTAGTGGAATTAGACGGCGAC<br>GTGAACGGTCATAAATTTAGCGTGCGCGGCGAGGGCGAAGGTGACG<br>CTACCAATGGTAAATTGACCCTGAAGTTTATTTGCACAACAGGCAAA<br>TTACCCGTTCCGTGGCCCACCTTAGTGACCACCCTGACCTATGGCGTT<br>CAGTGCTTCAGTCGTTACCCTAAACATATGAAACAACACGATTTTTC<br>AAATCAGCCATGCCTGAAGGATATGTTCAAGAGCGTACAATCAGCTT<br>CAAGAAGGATGGCACCTATAAAACGCGTGCGGAAGTGAAATTTGAA<br>GGCCGCACATTAGTAAATCGTATCGAACTGAAAGGTCGTGACTTCAA<br>AGAAGACGGCAACATTTTAGGCCATAAACTGGAATATAACTTTAATT<br>CTCATAACGTGTATATTACGGCCGATAAACAGAAGAATGGTATCAAG<br>GCAAATTTCAAAATTCGCCATAACGTGGAGGACGGCAGCGTTCAATT<br>AGCGGATCATTATCAACAAAACACGCCGATTGGTGATGGGCCTGTAC<br>TGTTACCTCGCAACCACTACCTGAGCACCCAATCTGCCCTGAGCAAA<br>GATCCGAAAGAAAAACGCGATCACATGGTTCTGTTAGAATTCGTGAC<br>CGCTGCAGGCATTACGCACGGAATGGACGAACGCTACAAGTAA |

| | |
|---|---|
| GFP(+6)-6Lys | ATGGGTCATCACCACCACCATCACGGTGGCGCTAGCAAAGGTGAAC<br>GTCTGTTTACTGGTGTAGTACCGATCTTAGTGGAATTAGACGGCGAC<br>GTGAACGGTCATAAATTTAGCGTGCGCGGCGAGGGCGAAGGTGACG<br>CTACCAATGGTAAATTGACCCTGAAGTTTATTTGCACAACAGGCAAA<br>TTACCCGTTCCGTGGCCCACCTTAGTGACCACCCTGACCTATGGCGTT<br>CAGTGCTTCAGTCGTTACCCTAAACATATGAAACAACACGATTTTTC<br>AAATCAGCCATGCCTGAAGGATATGTTCAAGAGCGTACAATCAGCTT<br>CAAGAAGGATGGCACCTATAAAACGCGTGCGGAAGTGAAATTTGAA<br>GGCCGCACATTAGTAAATCGTATCGAACTGAAAGGTCGTGACTTCAA<br>AGAAGACGGCAACATTTTAGGCCATAAACTGGAATATAACTTTAATT<br>CTCATAACGTGTATATTACGGCCGATAAACAGAAGAATGGTATCAAG<br>GCAAATTTCAAAATTCGCCATAACGTGGAGGACGGCAGCGTTCAATT<br>AGCGGATCATTATCAACAAAACACGCCGATTGGTGATGGGCCTGTAC<br>TGTTACCTCGCAACCACTACCTGAGCACCCAATCTGCCCTGAGCAAA<br>GATCCGAAAGAAAAACGCGATCACATGGTTCTGTTAGAATTCGTGAC<br>CGCTGCAGGCATTACGCACGGAATGGACGAACGCTACAAGAAAAAG<br>AAAAAGAAGAAATAA |
| R. Primer | 5'-TTTCTTCTTTTTCTTTTTCTTGTAGCGTTCGTCCATTCC-3' |
| F. Primer | 5'-<br>GAATGGACGAACGCTACAAGAAAAAGAAAAAGAAGAAATAATAAT<br>GAGGTACCCTCG-3' |
| GFP(+6)-6Arg | ATGGGTCATCACCACCACCATCACGGTGGCGCTAGCAAAGGTGAAC<br>GTCTGTTTACTGGTGTAGTACCGATCTTAGTGGAATTAGACGGCGAC<br>GTGAACGGTCATAAATTTAGCGTGCGCGGCGAGGGCGAAGGTGACG<br>CTACCAATGGTAAATTGACCCTGAAGTTTATTTGCACAACAGGCAAA<br>TTACCCGTTCCGTGGCCCACCTTAGTGACCACCCTGACCTATGGCGTT<br>CAGTGCTTCAGTCGTTACCCTAAACATATGAAACAACACGATTTTTC<br>AAATCAGCCATGCCTGAAGGATATGTTCAAGAGCGTACAATCAGCTT<br>CAAGAAGGATGGCACCTATAAAACGCGTGCGGAAGTGAAATTTGAA<br>GGCCGCACATTAGTAAATCGTATCGAACTGAAAGGTCGTGACTTCAA<br>AGAAGACGGCAACATTTTAGGCCATAAACTGGAATATAACTTTAATT<br>CTCATAACGTGTATATTACGGCCGATAAACAGAAGAATGGTATCAAG<br>GCAAATTTCAAAATTCGCCATAACGTGGAGGACGGCAGCGTTCAATT<br>AGCGGATCATTATCAACAAAACACGCCGATTGGTGATGGGCCTGTAC<br>TGTTACCTCGCAACCACTACCTGAGCACCCAATCTGCCCTGAGCAAA<br>GATCCGAAAGAAAAACGCGATCACATGGTTCTGTTAGAATTCGTGAC<br>CGCTGCAGGCATTACGCACGGAATGGACGAACGCTACAAGCGTCGC<br>CGGCGCCGTCGGTAA |
| R. Primer | 5'-CCGACGGCGCCGGCGACGCTTGTAGCGTTCGTCCATTCC-3' |

| | |
|---|---|
| F. Primer | 5'-<br>GAATGGACGAACGCTACAAGCGTCGCCGGCGCCGTCGGTAATAATG<br>AGGTACCCTCG-3' |
| GFP(+6)-<br>6Lys4Pepper | ATGGGTCATCACCACCACCATCACGGTGGCGCTAGCAAAGGTGAAC<br>GTCTGTTTACTGGTGTAGTACCGATCTTAGTGGAATTAGACGGCGAC<br>GTGAACGGTCATAAATTTAGCGTGCGCGGCGAGGGCGAAGGTGACG<br>CTACCAATGGTAAATTGACCCTGAAGTTTATTTGCACAACAGGCAAA<br>TTACCCGTTCCGTGGCCCACCTTAGTGACCACCCTGACCTATGGCGTT<br>CAGTGCTTCAGTCGTTACCCTAAACATATGAAACAACACGATTTTTC<br>AAATCAGCCATGCCTGAAGGATATGTTCAAGAGCGTACAATCAGCTT<br>CAAGAAGGATGGCACCTATAAAACGCGTGCGGAAGTGAAATTTGAA<br>GGCCGCACATTAGTAAATCGTATCGAACTGAAAGGTCGTGACTTCAA<br>AGAAGACGGCAACATTTTAGGCCATAAACTGGAATATAACTTTAATT<br>CTCATAACGTGTATATTACGGCCGATAAACAGAAGAATGGTATCAAG<br>GCAAATTTCAAAATTCGCCATAACGTGGAGGACGGCAGCGTTCAATT<br>AGCGGATCATTATCAACAAAACACGCCGATTGGTGATGGGCCTGTAC<br>TGTTACCTCGCAACCACTACCTGAGCACCCAATCTGCCCTGAGCAAA<br>GATCCGAAAGAAAAACGCGATCACATGGTTCTGTTAGAATTCGTGAC<br>CGCTGCAGGCATTACGCACGGAATGGACGAACGCTACAAGAAAAAG<br>AAAAAGAAGAaataataatgaggtaccctcgagtctggtaaagaaaccgctgctgcgaaatgcccacg<br>gaggatccccaatcgtggcgtgtcggcctctcccaatcgtggcgtgtcggcctctcccaatcgtggcgtgtcggcc<br>tctcccaatcgtggcgtgtcggcctct |
| GFP(+6)-<br>6Arg4Pepper | ATGGGTCATCACCACCACCATCACGGTGGCGCTAGCAAAGGTGAAC<br>GTCTGTTTACTGGTGTAGTACCGATCTTAGTGGAATTAGACGGCGAC<br>GTGAACGGTCATAAATTTAGCGTGCGCGGCGAGGGCGAAGGTGACG<br>CTACCAATGGTAAATTGACCCTGAAGTTTATTTGCACAACAGGCAAA<br>TTACCCGTTCCGTGGCCCACCTTAGTGACCACCCTGACCTATGGCGTT<br>CAGTGCTTCAGTCGTTACCCTAAACATATGAAACAACACGATTTTTC<br>AAATCAGCCATGCCTGAAGGATATGTTCAAGAGCGTACAATCAGCTT<br>CAAGAAGGATGGCACCTATAAAACGCGTGCGGAAGTGAAATTTGAA<br>GGCCGCACATTAGTAAATCGTATCGAACTGAAAGGTCGTGACTTCAA<br>AGAAGACGGCAACATTTTAGGCCATAAACTGGAATATAACTTTAATT<br>CTCATAACGTGTATATTACGGCCGATAAACAGAAGAATGGTATCAAG<br>GCAAATTTCAAAATTCGCCATAACGTGGAGGACGGCAGCGTTCAATT<br>AGCGGATCATTATCAACAAAACACGCCGATTGGTGATGGGCCTGTAC<br>TGTTACCTCGCAACCACTACCTGAGCACCCAATCTGCCCTGAGCAAA<br>GATCCGAAAGAAAAACGCGATCACATGGTTCTGTTAGAATTCGTGAC<br>CGCTGCAGGCATTACGCACGGAATGGACGAACGCTACAAGCGTCGC<br>CGGCGCCGTCGGTaataatgaggtaccctcgagtctggtaaagaaaccgctgctgcgaaatgcccac<br>ggaggatccccaatcgtggcgtgtcggcctctcccaatcgtggcgtgtcggcctctcccaatcgtggcgtgtcggc<br>ctctcccaatcgtggcgtgtcggcctct |
| 4Pepper<br>aptamer | CCCAATCGTGGCGTGTCGGCCTCTCCCAATCGTGGCGTGTCGGCCTCT<br>CCCAATCGTGGCGTGTCGGCCTCTCCCAATCGTGGCGTGTCGGCCTCT |

Table 2. Protein Sequences
***Bold** residues correspond to those mutated from the sfGFP sequence

| | |
|---|---|
| sfGFP | MGHHHHHHGGASKGEELFTGVVPILVELDGDVNGHKFSVRGEGEGDA<br>TNGKLTLKFICTTGKLPVPWPTLVTTLTYGVQCFSRYPDHMKQHDFFKS<br>AMPEGYVQERTISFKDDGTYKTRAEVKFEGDTLVNRIELKGIDFKEDGN<br>ILGHKLEYNFNSHNVYITADKQKNGIKANFKIRHNVEDGSVQLADHYQ<br>QNTPIGDGPVLLPDNHYLSTQSALSKDPNEKRDHMVLLEFVTAAGITHG<br>MDELYK* |
| GFP(0) | MGHHHHHHGGASKGE**R**LFTGVVPILVELDGDVNGHKFSVRGEGEGDA<br>TNGKLTLKFICTTGKLPVPWPTLVTTLTYGVQCFSRYPDHMKQHDFFKS<br>AMPEGYVQERTISFKDDGTYKTRAEVKFEGDTLVNRIELKG**R**DFKEDG<br>NILGHKLEYNFNSHNVYITADKQKNGIKANFKIRHNVEDGSVQLADHY<br>QQNTPIGDGPVLLP**R**NHYLSTQSALSKDP**K**EKRDHMVLLEFVTAAGITH<br>GMDE**R**YK* |
| GFP(0)-6Lys | MGHHHHHHGGASKGE**R**LFTGVVPILVELDGDVNGHKFSVRGEGEGDA<br>TNGKLTLKFICTTGKLPVPWPTLVTTLTYGVQCFSRYPDHMKQHDFFKS<br>AMPEGYVQERTISFKDDGTYKTRAEVKFEGDTLVNRIELKG**R**DFKEDG<br>NILGHKLEYNFNSHNVYITADKQKNGIKANFKIRHNVEDGSVQLADHY<br>QQNTPIGDGPVLLP**R**NHYLSTQSALSKDP**K**EKRDHMVLLEFVTAAGITH<br>GMDE**R**YK**KKKKKK*** |
| GFP(0)-6Arg | MGHHHHHHGGASKGE**R**LFTGVVPILVELDGDVNGHKFSVRGEGEGDA<br>TNGKLTLKFICTTGKLPVPWPTLVTTLTYGVQCFSRYPDHMKQHDFFKS<br>AMPEGYVQERTISFKDDGTYKTRAEVKFEGDTLVNRIELKG**R**DFKEDG<br>NILGHKLEYNFNSHNVYITADKQKNGIKANFKIRHNVEDGSVQLADHY<br>QQNTPIGDGPVLLP**R**NHYLSTQSALSKDP**K**EKRDHMVLLEFVTAAGITH<br>GMDE**R**YK**RRRRRR*** |
| GFP(+6) | MGHHHHHHGGASKGE**R**LFTGVVPILVELDGDVNGHKFSVRGEGEGDA<br>TNGKLTLKFICTTGKLPVPWPTLVTTLTYGVQCFSRYP**K**HMKQHDFFKS<br>AMPEGYVQERTISFK**K**DGTYKTRAEVKFEG**R**TLVNRIELKG**R**DFKEDG<br>NILGHKLEYNFNSHNVYITADKQKNGIKANFKIRHNVEDGSVQLADHY<br>QQNTPIGDGPVLLP**R**NHYLSTQSALSKDP**K**EKRDHMVLLEFVTAAGITH<br>GMDE**R**YK* |
| GFP(+6)-6Lys | MGHHHHHHGGASKGE**R**LFTGVVPILVELDGDVNGHKFSVRGEGEGDA<br>TNGKLTLKFICTTGKLPVPWPTLVTTLTYGVQCFSRYP**K**HMKQHDFFKS<br>AMPEGYVQERTISFK**K**DGTYKTRAEVKFEG**R**TLVNRIELKG**R**DFKEDG<br>NILGHKLEYNFNSHNVYITADKQKNGIKANFKIRHNVEDGSVQLADHY |

| | |
|---|---|
| | QQNTPIGDGPVLLP**R**NHYLSTQSALSKDP**K**EKRDHMVLLEFVTAAGITH GMDE**R**YK**KKKKKK***|
| GFP(+6)-6Arg | MGHHHHHHGGASKGE**R**LFTGVVPILVELDGDVNGHKFSVRGEGEGDA TNGKLTLKFICTTGKLPVPWPTLVTTLTYGVQCFSRYP**K**HMKQHDFFKS AMPEGYVQERTISFK**K**DGTYKTRAEVKFEG**R**TLVNRIELKG**R**DFKEDG NILGHKLEYNFNSHNVYITADKQKNGIKANFKIRHNVEDGSVQLADHY QQNTPIGDGPVLLP**R**NHYLSTQSALSKDP**K**EKRDHMVLLEFVTAAGITH GMDE**R**YK**RRRRRR***|

### Section 2: Supplemental Figures

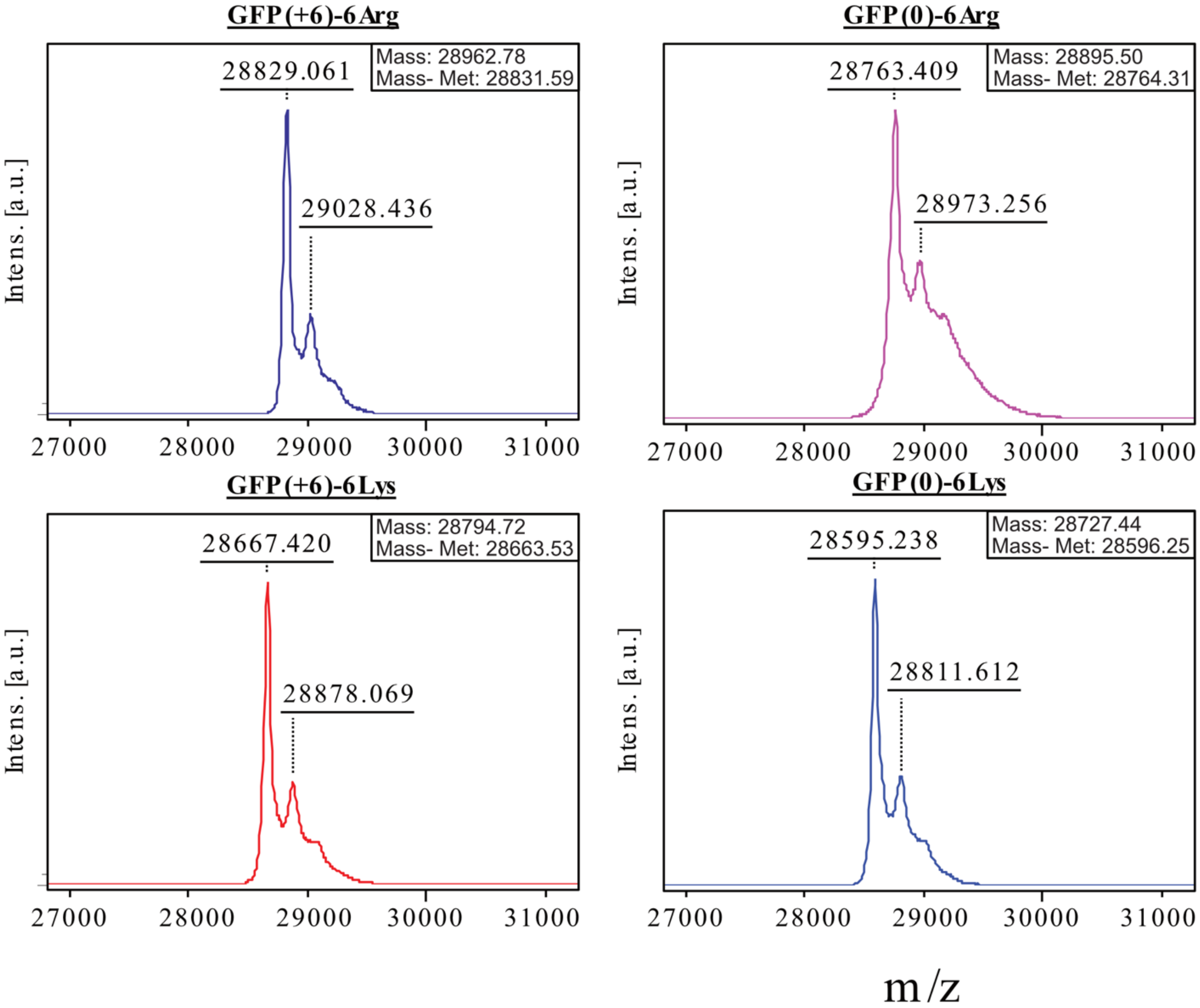


**Supplemental Figure 1.** MALDI-TOF MS of purified GFP(0)-6Arg, GFP(0)-6Lys, GFP(+6)-6Arg, and GFP(+6)-6Lys. All samples were analyzed on a Bruker Ultraflextreme MALDI-TOF/TOF on positive mode, in a matrix of 30% acetonitrile in water with 0.1% TFA and sinapinic acid to saturation.

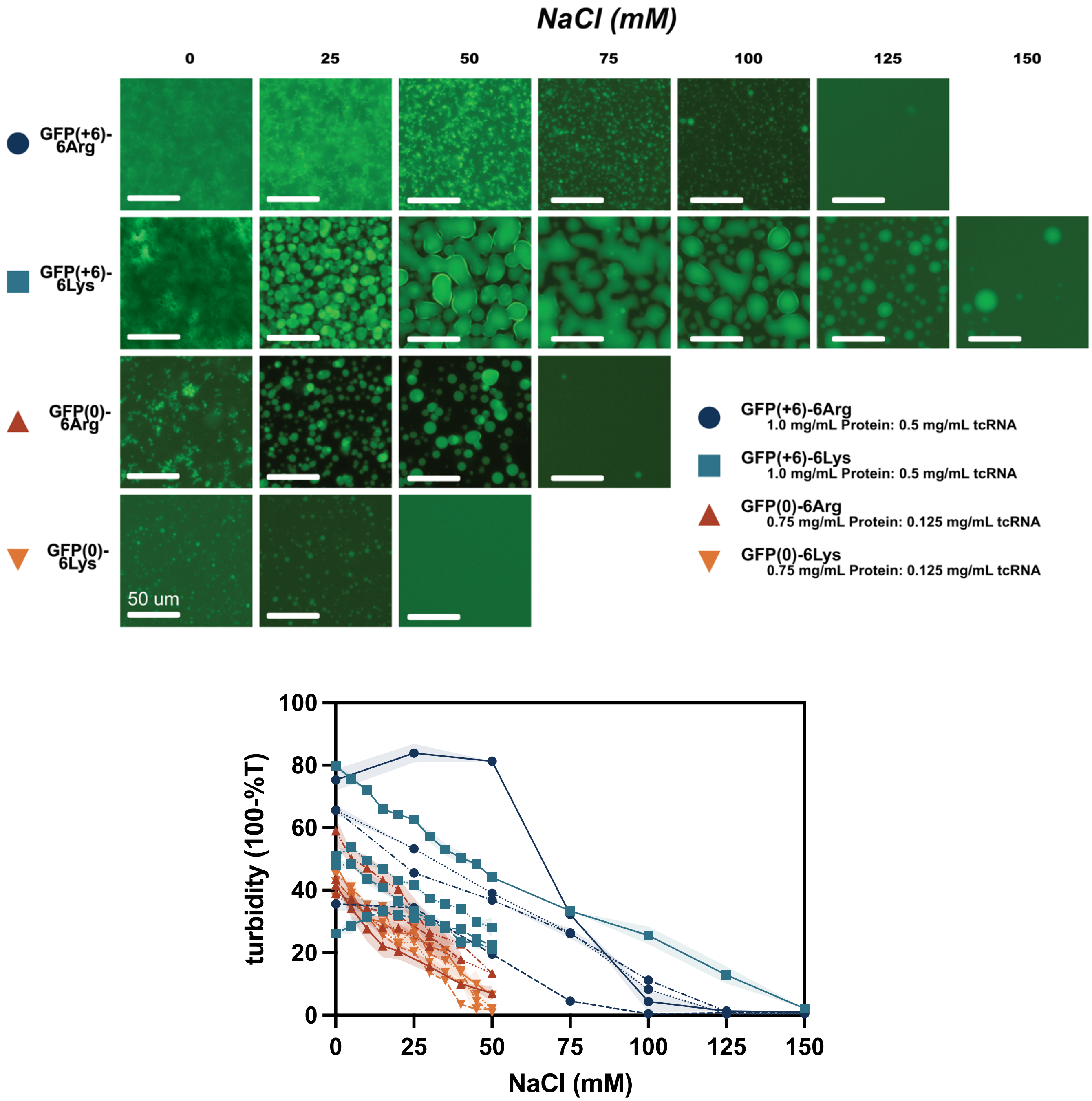


**<u>Supplementary Figure 2</u>**. (A) Microscopy images of *in vitro* protein:RNA condensates with increasing concentrations of NaCl. Protein-to-RNA ratios were selected based on the peak turbidity in a 2D phase diagram. All experiments performed in a pH 7.4, 10 mM Tris buffer. All images were collected on a 20X/ 0.4 NA, LPlanFL PH2 objective (Olympus) on GFP $\lambda_{ex}$ = 470-522 nm; $\lambda_{em}$ = 525-550 nm; EVOS GFP light cube) using an EVOS FL AUTO 2 inverted fluorescent microscope. (B) NaCl titration experiments with varying concentrations of the engineered protein variants and tcRNA. GFP(+6)-6Arg (dark blue): solid line: 1 mg/mL protein with 0.5 mg/mL tcRNA; mixed dash and dots line: 1 mg/mL protein with 0.125 mg/mL tcRNA; dotted line: 0.5 mg/mL protein with 0.125 mg/mL tcRNA; dashed line: 0.25 mg/mL protein with

0.125 mg/mL tcRNA; GFP(+6)-6Lys (light blue): solid line: 1 mg/mL protein with 0.5 mg/mL tcRNA; mixed dash and dots line: 1 mg/mL protein with 0.125 mg/mL tcRNA; dotted line: 0.5 mg/mL protein with 0.125 mg/mL tcRNA; dashed line: 0.25 mg/mL protein with 0.125 mg/mL tcRNA; GFP(0)-6Arg (dark orange): solid line: 1 mg/mL protein with 0.125 mg/mL tcRNA; mixed dash and dots line: 0.75 mg/mL protein with 0.125 mg/mL tcRNA; dotted line: 0.75 mg/mL protein with 0.0625 mg/mL tcRNA; dashed line: 0.5 mg/mL protein with 0.0625 mg/mL tcRNA; GFP(0)-6Lys (light orange): solid line: 1 mg/mL protein with 0.125 mg/mL tcRNA; mixed dash and dots line: 0.75 mg/mL protein with 0.125 mg/mL tcRNA; dotted line: 0.75 mg/mL protein with 0.0625 mg/mL tcRNA; dashed line: 0.5 mg/mL protein with 0.0625 mg/mL tcRNA.

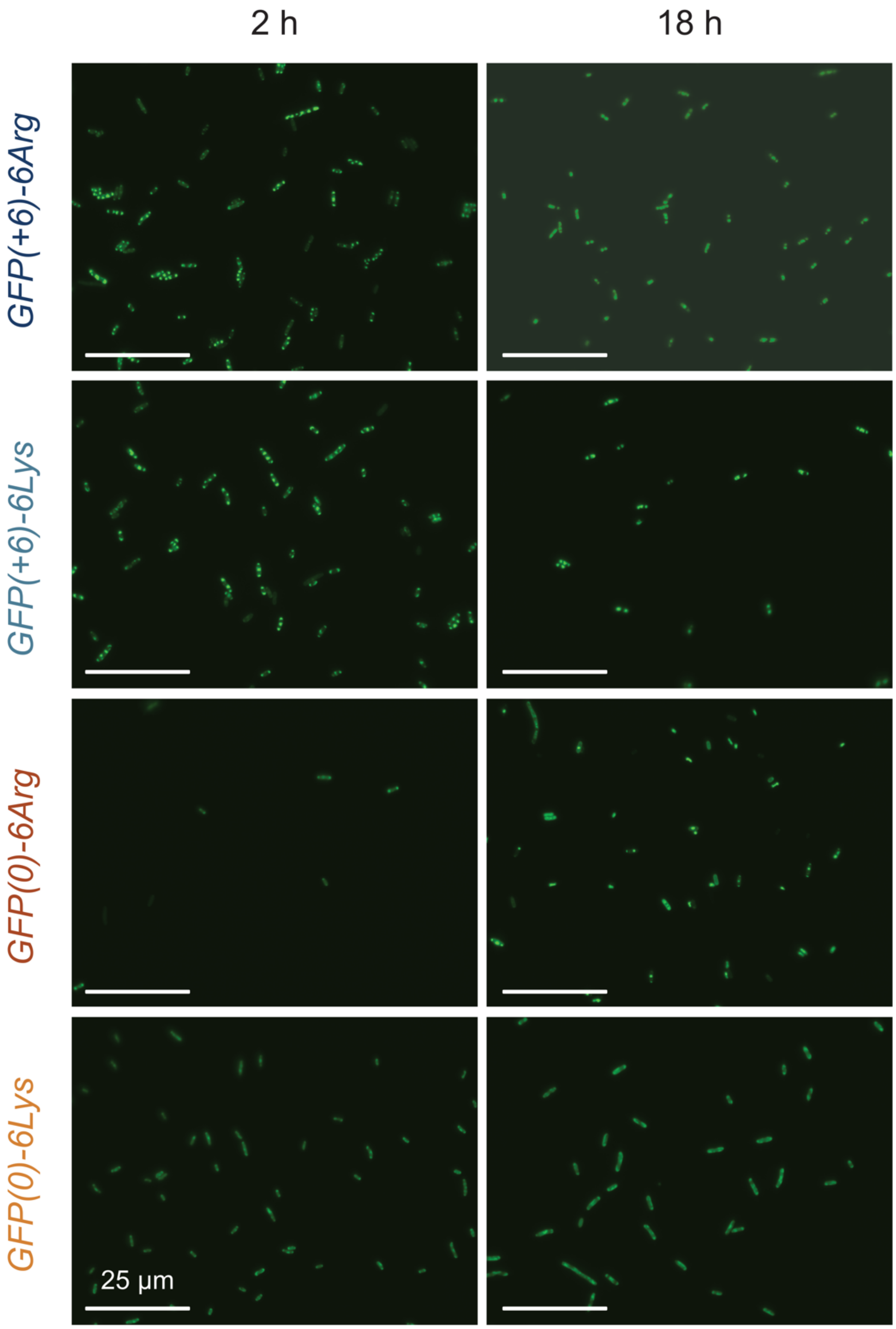


**<u>Supplemental Figure 3</u>.** Uncropped microscopy images of GFP(0)-6Arg, GFP(0)-6Lys, GFP(+6)-6Arg and GFP(+6)-6Lys proteins grown in NiCo expression cells. All images were collected on a 100X oil 1.40 NA UPlanSApo objective (Olympus) on GFP $\lambda_{ex}$ = 470-522 nm; $\lambda_{em}$ = 525-550 nm; EVOS GFP light cube) using an EVOS FL AUTO 2 inverted fluorescent microscope.

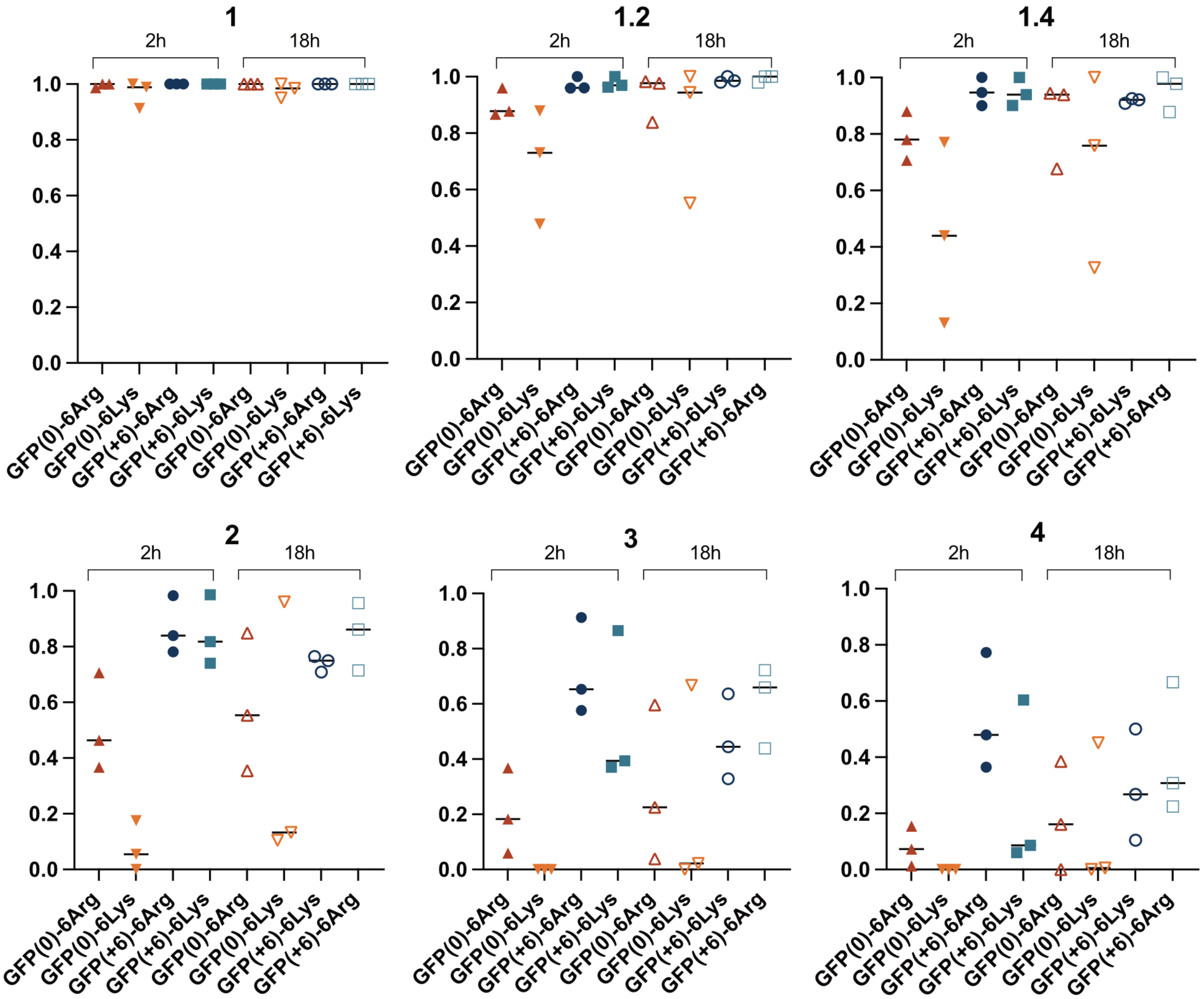


| Strain | Condensate-to-cytoplasm ratio | |
|---|---|---|
| | **2 h** | **18 h** |
| GFP(0) | 1.2 ± 0.3 | 1.2 ± 0.1 |
| GFP(0)-6Lys | 1.6 ± 0.3 | 1.9 ± 0.6 |
| GFP(0)-6Arg | 2.7 ± 0.8 | 3.1 ± 1.1 |
| GFP(+6) | 2.4 ± 0.6 | 2.6 ± 1.3 |
| GFP(+6)-6Lys | 4.0 ± 1.0 | 3.1 ± 1.0 |
| GFP(+6)-6Arg | 5.2 ± 2.2 | 4.1 ± 1.6 |

**Supplementary Figure 4.** (A) Fraction of cells with condensates. Microscopy images were analyzed using the MicrobeJ FIJI plugin to identify cells and a custom MATLAB script (available on Github) was used to identify condensates. The numbers above each graph panel corresponds to the threshold ratio to identify a condensate from the cytoplasm. (B) The estimated condensate-to-cytoplasm ratio for each strain at 2 and 18 h post-induction was averaged across all cells in three biological replicates, shown are the average and standard deviation of this ratio for each strain (data shown in Figure 3B).

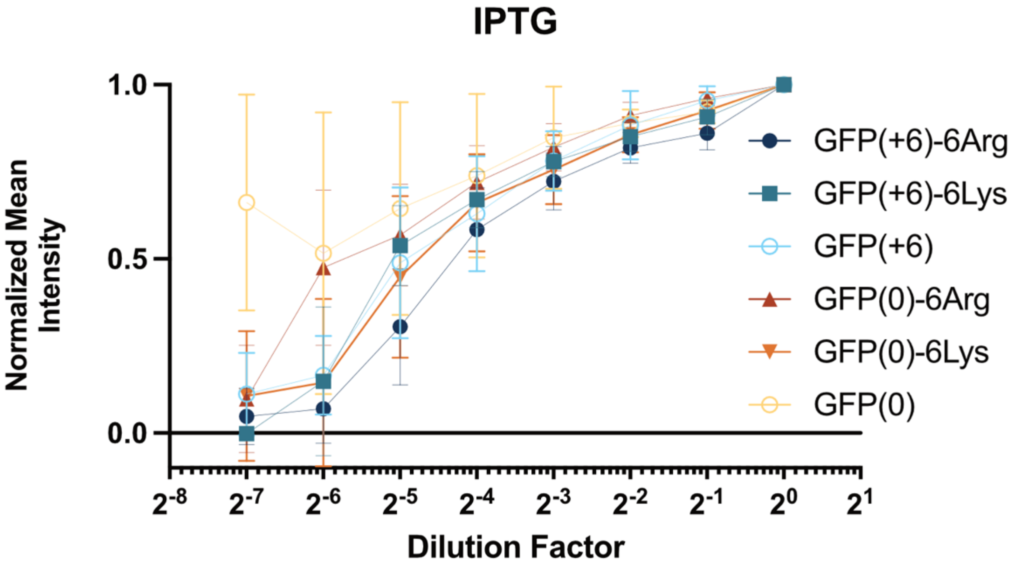


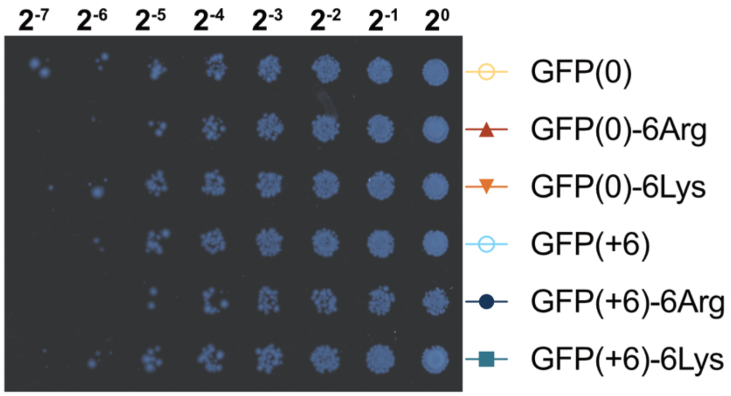


**Supplementary Figure 5.** Spotting assay on LB agar plates. Bacterial survival on a plate supplemented with both ampicillin and 1 mM IPTG to induce protein expression is correlated to condensate formation. Higher charge proteins and arginine-tagged proteins result in lower cell growth.

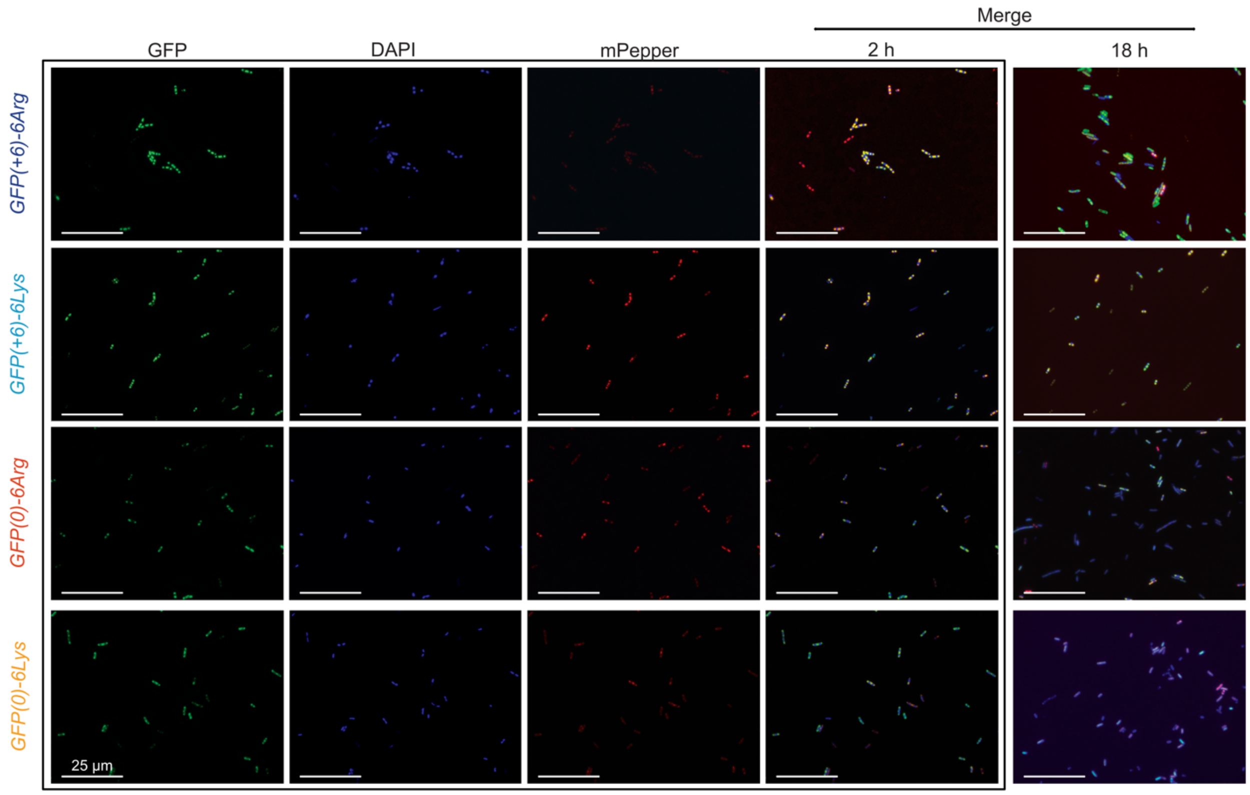


**<u>Supplemental Figure 6</u>.** Uncropped microscopy images of GFP(0)-6Arg, GFP(0)-6Lys, GFP(+6)-6Arg and GFP(+6)-6Lys proteins grown in NiCo expression cells with DAPI and HBC620 fluorescent dyes to stain DNA and RNA, respectively. All images were collected on a 100X oil 1.40 NA UPlanSApo objective (Olympus) on GFP ($\lambda_{ex}$ = 470-522 nm; $\lambda_{em}$= 525-550 nm; EVOS GFP light cube), DAPI (Thermo-Fisher 62248, ($\lambda_{ex}$= 360 nm; $\lambda_{em}$= 460 nm); EVOS DAPI light cube) and HBC620 (FD Biotech Cat No. H16201, ($\lambda_{ex}$ 570 nm, $\lambda_{em}$= 620 nm); EVOS Texas Red light cube) using an EVOS FL AUTO 2 inverted fluorescent microscope.

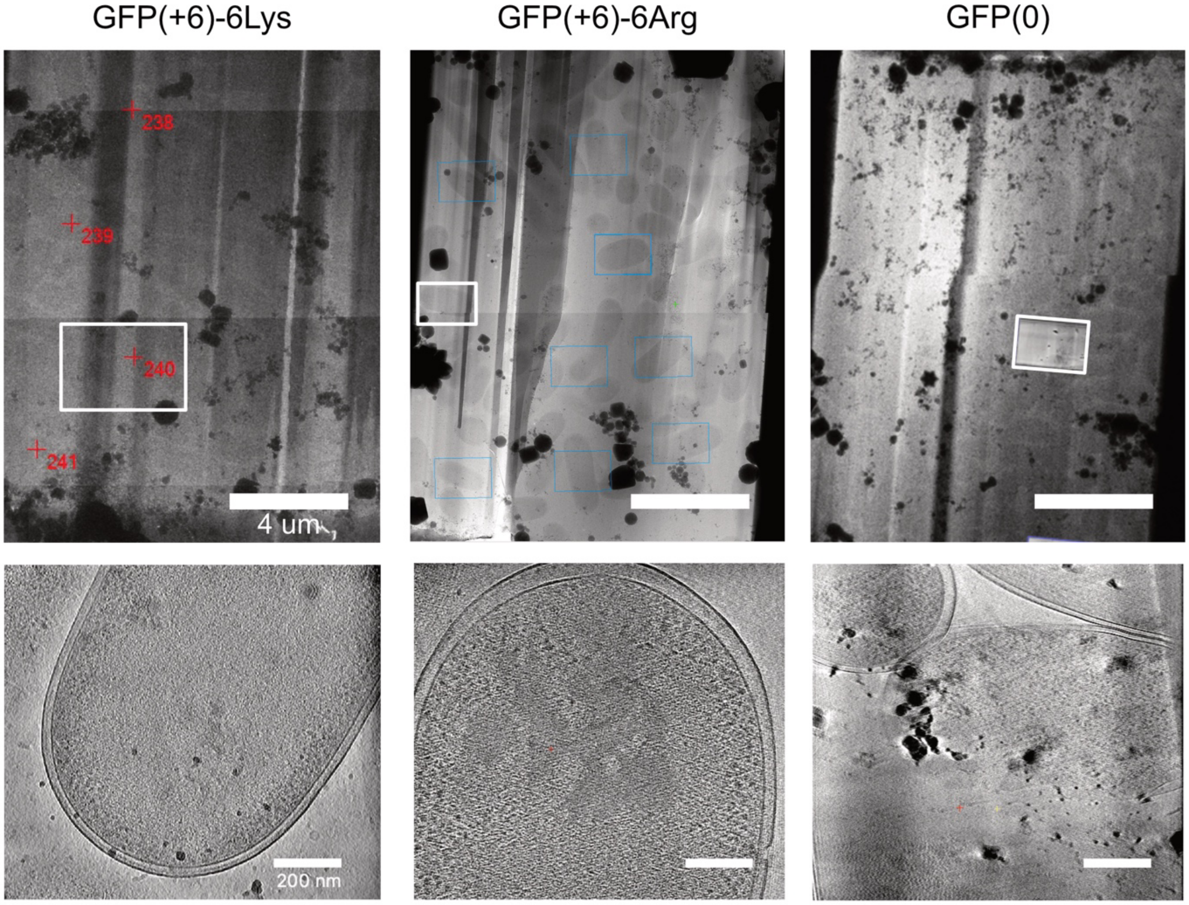


**Supplementary Figure 7.** Cryo-CLEM lamella (top) and tomograms (bottom) for GFP(+6)-6Lys (left), GFP(+6)-6Arg (middle), and GFP(0) (right) expressed in NiCo cells. White box indicates the region of the lamella shown in the tomogram at bottom
